# AlphaFold3 and Intrinsically Disordered Proteins: Reliable Monomer Prediction, Unpredictable Multimer Performance

**DOI:** 10.64898/2025.12.05.691730

**Authors:** Tuan Minh Dao, Sebastien Ghent, Vladimir N. Uversky, Taseef Rahman

## Abstract

AlphaFold3 represents a major advance in protein structure prediction, yet its performance on intrinsically disordered proteins remains uncharacterized. We present the first systematic evaluation of AF3 on disordered systems, revealing a striking dichotomy. For monomers, AF3’s pLDDT scores reliably predict disorder (MCC: 0.693), matching AlphaFold2 and rivaling dedicated predictors. This consistency across fundamentally different architectures confirms that disorder prediction emerges from training data, not model design. For multimers, the picture grows complex. Despite comparable aggregate performance (mean DockQ: 0.563 vs 0.571), AF3 and AF2 achieve these results through fundamentally different mechanisms. Conventional structural features explain 58% of AF2’s variance but only 42% of AF3’s. Users cannot predict when AF3 will succeed or fail from interface properties alone. On disorder-to-order transitions (MFIB benchmark), both models perform equally well, successfully predicting final folded states. Yet seed variance analysis reveals that AF3 rarely converges reliably as most cases show substantial variance across independent runs, with occasional high-scoring outlier seeds within an otherwise low-scoring distribution, indicating the model can sometimes locate a correct solution without consistently identifying it. Our findings establish AF3 as reliable for the prediction of monomer disorder but unpredictable for multimers. Architectural innovation alone cannot overcome training data bias. Progress demands disorder-enriched datasets and ensemble sampling, not merely novel architectures.

## Introduction

Intrinsically disordered regions (IDRs) constitute 30-40% of eukaryotic proteomes and orchestrate essential cellular processes, including regulation, signaling, and biomolecular condensate formation Xue et al. (2012); Holehouse et al. (2017); Wright and Dyson (2015); Miao and Chong (2025); Qin et al. (2025). Proteins consisting wholly or predominantly of such regions are termed intrinsically disordered proteins (IDPs) throughout this manuscript. Unlike well-folded proteins, IDRs lack a stable three-dimensional structure yet enable diverse functional roles through structural plasticity. Their dysregulation drives pathologies from cancer to neurodegeneration, making their characterization a therapeutic imperative. The challenge intensifies at protein-protein interfaces, where 15-40% involve intrinsic disorder Tompa and Fuxreiter (2008). IDRs mediate binding through diverse mechanisms—disorder-to-order transitions, fuzzy complexes that maintain dynamic disorder while bound, or fly-casting mechanisms that use conformational dynamics for target recognition Dyson and Wright (2005); Sugase et al. (2007). Understanding how computational models handle these dynamic, disordered interfaces is essential for interaction network analysis and structure-based drug design.

AlphaFold2 (AF2)’s 2021 debut revolutionized structural biology, achieving near-experimental accuracy for folded proteins Jumper et al. (2021). AlphaFold3 (AF3) Abramson et al. (2024) emerged in 2024 with architectural reinvention—replacing the evoformer with diffusion-based generation, implementing unified molecular representations, and expanding scope to diverse biomolecular complexes. Yet amid these advances, an intriguing observation about AF2 went largely unexplored in AF3: AF2, never explicitly trained for disorder prediction, proved remarkably competent at identifying disordered regions. Studies revealed that AF2’s confidence metric, pLDDT, correlates strongly with disorder propensity Wilson et al. (2022), matching or exceeding specialized disorder predictors. This emergent capability raised a fundamental question—would AF3’s architectural evolution preserve this proficiency?

Despite AF3’s prominence, **no systematic disorder analysis exists**. This gap is particularly striking for disorder-mediated interactions. While AlphaFold2-Multimer, the multimer-prediction pipeline built on AF2, has been shown to accurately capture interactions involving IDRs Omidi et al. (2024), AF3’s performance on disorder-containing complexes remains uncharacterized. The transition to diffusion-based generation raises critical questions: Does diffusion sampling better explore conformational ensembles, or does it inappropriately collapse diverse states into single structures? Can AF3 handle fuzzy complexes where partners remain dynamically disordered while bound? Without systematic benchmarking, users lack guidance on when to trust AF3 predictions for the substantial fraction of biological interactions involving disordered regions.

We present the first comprehensive evaluation of AlphaFold3’s performance on disordered systems, revealing both expected strengths and surprising limitations. For monomers, we leverage DisProt-PDB with experimentally validated disorder annotations to evaluate classification accuracy and residue-level propensity Wilson et al. (2022). For multimers, we constructed a benchmark dataset of 90 CAPRI complexes deposited after April 2021, ensuring no training contamination, and systematically stratified predictions by buried surface area, interface coil content, and other structural features to identify determinants of success and failure. We further evaluated AF3 on 90 fuzzy complexes from the MFIB benchmark to assess performance on the most challenging class of disorder-mediated interactions.

Our findings reveal nuanced complexity beneath apparent simplicity. For monomers, AF3 matches AF2 despite architectural differences, with both rivaling dedicated disorder predictors and validating pLDDT as a reliable disorder indicator. However, extending to multimeric assemblies uncovers unexpected complexity. While AF3 generally performs comparably to AF2 with similar mean DockQ scores, aggregate statistics mask dramatic individual divergence. A subset of multimers shows striking quality differences that conventional structural features cannot fully explain. Systematic analysis across our data set reveals that the interface pLDDT and secondary structure metrics account for only half of the observed DockQ variation, suggesting that AF3 employs fundamentally different mechanisms for multimer assembly. On fuzzy complexes, both models achieve comparable performance, demonstrating that AF3’s architectural innovations neither substantially improve nor worsen the prediction of genuinely dynamic, disordered interfaces. These results have important implications for practitioners using AF3 and highlight critical gaps in our understanding of how modern structure prediction models handle disorder.

## Methods

### Datasets

We evaluated monomer disorder prediction using the DisProt-PDB dataset from Wilson et al. Wilson et al. (2022), derived from DisProt database release 2018_11. This dataset comprises 478 protein targets with experimentally validated disorder annotations. Residues annotated as disordered in DisProt received label 1, while residues with ordered structures in the Protein Data Bank (PDB) received label 0. When annotations conflicted, disorder annotations took precedence. Residues lacking experimental evidence from either source were masked and excluded from analysis, ensuring that the DisProt-PDB dataset contains only experimentally validated residues and provides a gold-standard benchmark for disorder prediction evaluation.

For multimer analysis, we constructed Benchmark-90 (B90) from the blind prediction dataset of Omidi et al. Omidi et al. (2024). The parent dataset contains 210 nonredundant IDR-receptor dimers from PDB entries released after 2018-04-30, curated through stringent criteria including binary protein complexes with resolution *≤* 3.5Å or NMR (Nuclear Magnetic Resonance) structures, IDR-receptor classification via radius-of-gyration scaling requiring interchain contacts *≤* 5Å, removal of disulfide-rich complexes, and minimum sequence lengths of 20 residues for IDRs and 70 residues for receptors. To ensure zero training contamination for AF3 evaluation, we selected only complexes deposited after September 30, 2021, yielding 97 eligible structures. From these, we curated 90 complexes to achieve an approximately normal distribution of AF2 DockQ scores, providing balanced representation across prediction difficulty levels. The dataset maintains sequence nonredundancy through clustering at 30% identity and 80% coverage for both IDR and receptor chains.

We additionally evaluated disorder-to-order transitions using 90 complexes from the Mutual Folding-Induced Binding (MFIB) 2.0 database Fichó et al. (2025), which catalogs protein complexes where IDRs fold upon binding to their partners. This benchmark represents the most challenging class of disorder-mediated interactions, where IDRs adopt well-defined structures only in the presence of their binding partners.

### Structure Prediction and Analysis

All 90 B90 complexes were predicted using both AlphaFold3 Abramson et al. (2024) via the AlphaFold Server and AlphaFold2-Multimer v3 Jumper et al. (2021) with default parameters. For each complex, we submitted the protein sequences and extracted the top-ranked predictions along with associated confidence metrics, including pLDDT (per-residue confidence), pTM (predicted template modeling score), iPTM (interface predicted template modeling score), and PAE (predicted aligned error). These metrics provide complementary views of model confidence at both the residue and complex levels. The use of both models allows us to assess whether AF3’s architectural innovations translate to improved performance on disorder-containing complexes.

To determine whether AF3’s performance variation arises from stochastic sampling or systematic failures, we selected six cases exhibiting significant AF2-AF3 performance divergence and generated five additional AF3 predictions for each using different random seeds. This analysis reveals whether the diffusion-based architecture produces consistent predictions across independent runs or whether performance variability reflects genuine sampling diversity.

The predicted structures were aligned to native PDB chains using PyMOL Schrödinger, LLC (2015), and all subsequent analyses were restricted to experimentally resolved residues to ensure a fair evaluation. Interface residues were defined using a 5Å distance cutoff between chains, a standard threshold that captures both direct contacts and near-neighbor interactions relevant to binding. We assessed interface prediction accuracy using DockQ v2 Mirabello and Wallner (2024), a comprehensive metric that ranges from 0 (incorrect) to 1 (native-like) and integrates interface RMSD, ligand RMSD, and fraction of native contacts. DockQ provides a single, interpretable score that captures multiple aspects of complex quality, making it ideal for systematic benchmarking. We further categorized predictions according to CAPRI quality standards as Incorrect (DockQ *<* 0.23), Acceptable (0.23–0.49), Medium (0.49–0.80), and High (*>* 0.80), enabling comparison with previous blind prediction assessments and providing intuitive quality thresholds.

Secondary structure assignments were performed using DSSP (Dictionary of Secondary Structure of Proteins) Hekkelman et al. (2025) for both predicted and native structures. DSSP matching accuracy was calculated as the fraction of interface residues with matching secondary structure assignments (helix, sheet, or coil) between predicted and native structures. This metric provides insight into whether models correctly predict local conformational preferences at binding interfaces, which is particularly relevant for IDRs that may adopt structured conformations upon binding. We characterize multiple additional interface properties to understand their relationship with prediction quality. Buried surface area (BSA) was calculated using PyMOL to quantify interface size, with larger interfaces typically providing more structural constraints for prediction. The disorder content was calculated as the fraction of the interface residues predicted as disordered, using optimized thresholds of pLDDT *<* 70 for AF3 and pLDDT *<* 68 for AF2 (Tables S1, S2, and S3). Interface hydrophobicity was characterized using the Kyte-Doolittle scale Kyte and Doolittle (1982), which ranges from hydrophilic (negative values) to hydrophobic (positive values). For contact accuracy analysis, we calculated recall as the ratio of true contacts predicted to total true contacts, and identified over-contacted complexes where predicted contacts exceeded true contacts, a common failure mode in disorder-containing interfaces.

### Disorder Prediction Evaluation

For the prediction of monomer disorder, we determined optimal pLDDT thresholds by systematically testing values from 60 to 80 in increments of 2, following the methodology of Wilson et al. Wilson et al. (2022). The threshold maximizing Matthews Correlation Coefficient (MCC) was selected for each model, yielding optimal values of 68 for AF2, 70 for AF3, and 74 for ESMFold. These model-specific thresholds account for differences in confidence score calibration across architectures.

We computed a comprehensive set of metrics to evaluate disorder prediction performance. The Matthews Correlation Coefficient (MCC) served as our primary metric for classification accuracy, as it provides a balanced measure even with imbalanced class distributions. We calculated the Area Under ROC Curve (AUC) to assess discriminative power across all possible thresholds, providing a threshold-independent performance measure. F-maximum (F-max) was computed as the maximum F1 score across the precision-recall curve, emphasizing performance at optimal operating points. Root Mean Square Deviation (RMSD) quantified the difference between predicted and experimental disorder fractions at the protein level. We also calculated True Positive Rate (TPR, sensitivity for disordered residues) and True Negative Rate (TNR, specificity for ordered residues) to assess model performance on each class separately.

We calculated per-amino-acid disorder frequencies for ground truth annotations and all model predictions to assess whether models capture sequence-encoded disorder determinants. Pearson correlation coefficients were computed between model predictions and experimental values, with values near 1.0 indicating accurate recapitulation of amino acid disorder preferences. This analysis reveals whether models learn genuine biophysical principles or merely reflect overall disorder tendencies.

For monomer disorder prediction, we compared AlphaFold2 and AlphaFold3 Jumper et al. (2021); Abramson et al. (2024) pLDDT-based predictions against 11 established disorder predictors spanning multiple methodological approaches. Specialized predictors included spot_disorder1, spot_disorder2, aucpred, spot_disorder_s, and predisorder, which were explicitly trained for disorder prediction. We included RawMSA as a representative MSA-based method that leverages evolutionary information. Sequence-based predictors included fidpnn, disomine, espritz_d, fidplr, and aucpred_np, which rely solely on amino acid sequence features. Finally, we compared against ESMFold pLDDT as a representative language model-based approach. All predictors were evaluated using the same DisProt-PDB dataset and metrics to ensure fair comparison, allowing us to contextualize structure prediction model performance relative to dedicated disorder prediction tools.

### Regression and Statistical Analysis

To understand which conventional structural features determine prediction quality, we constructed linear regression models relating interface properties to DockQ scores. We first built univariate models with disorder fraction or secondary structure matching (DSSP accuracy) as sole predictors, then constructed a multivariate model combining both features to assess their joint explanatory power:

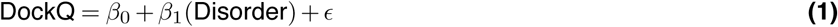

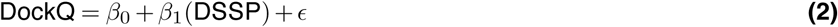

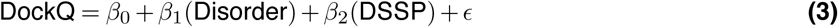

where Disorder represents interface disorder fraction and DSSP represents secondary structure matching accuracy. We computed *R*^2^ values to quantify the proportion of variance explained by each model and calculated incremental *R*^2^ (Δ*R*^2^) to assess relative feature importance. Specifically, we compared *R*^2^ values when adding DSSP to a disorder-only model versus adding disorder to a DSSP-only model, revealing which feature provides greater explanatory power. This analysis tests whether AF3’s diffusion-based architecture introduces dependencies beyond those captured by simple linear relationships.

We employed two-tailed Mann-Whitney U tests to compare AF2 and AF3 performance metrics, as these nonparametric tests make no distributional assumptions and are robust to outliers. We set a significance threshold of *α* = 0.05 for all tests. When performing multiple comparisons across disorder categories (low, medium, high), we applied the Bonferroni correction to control the family-wise error rate. For continuous variables, we computed Pearson correlation coefficients to quantify linear relationships between interface properties and prediction quality, including disorder versus DockQ and DSSP accuracy versus DockQ. Correlation strength was interpreted as weak (*|r| <* 0.3), moderate (0.3 *≤ |r| <* 0.7), or strong (*|r| ≥* 0.7).

### Molecular Recognition Feature Analysis

For the MFIB benchmark, we analyzed the models’ ability to identify Molecular Recognition Features (MoRFs), which are short disordered segments that fold upon binding. We first computed delta pLDDT as the difference between bound and unbound predictions for MoRF regions, hypothesizing that successful MoRF predictors should show reduced confidence in these regions. We then calculated direct prediction accuracy by applying optimal disorder thresholds to identify MoRFs prospectively. Finally, we correlated Mean Confidence Weighted (MCW) scores with average pLDDT values to assess whether confidence metrics contain information about binding-induced folding propensity. This analysis reveals whether models can prospectively identify which disordered regions will undergo folding transitions, a critical capability for understanding protein interaction networks.

### Software and Data Availability

Structure predictions were generated using the AlphaFold3 Server for AF3 predictions (alphafoldserver. com/). and AlphaFold2-Multimer v3 (sokrypton/ColabFold/blob/main/AlphaFold2.ipynb) for AF2 predictions. Structural analyses were performed using PyMOL Schrödinger, LLC (2015) for structure alignment, visualization, and buried surface area calculations, and DSSP Hekkelman et al. (2025) for secondary structure assignment. Interface quality was assessed using DockQ v2 Mirabello and Wallner (2024). Statistical analyses and data processing were performed using Python with the scipy library for statistical tests, numpy for numerical computations, and pandas for data manipulation. Visualizations were generated using matplotlib for plotting and seaborn for statistical graphics. The DisProt-PDB dataset used for monomer disorder evaluation is publicly available from Wilson et al. Wilson et al. (2022). The B90 benchmark dataset, including PDB identifiers and curation criteria, along with all predicted structures and computed metrics have been made available at github.com/Daominh09/AlphaFold3_IDP_Benchmark.

## Results

### AlphaFold3 pLDDT is a reliable predictor of disorder

To evaluate AlphaFold3’s per-residue confidence metric (pLDDT) as a disorder predictor, we benchmarked it against 11 established methods using the DisProt-PDB dataset. Our analysis reveals that pLDDT-based predictions consistently rank among the top-performing disorder predictors across multiple metrics. The optimal threshold for pLDDT was determined (see tables S1,S2, and S3) following the method described in Wilson et al. (2022). pLDDT from AlphaFold2 achieved the highest Matthews Correlation Coefficient (MCC) of 0.701, with AlphaFold3 close behind at 0.693, outperforming all sequence-based predictors (Figure S2). Three of the top five methods utilize pLDDT scores (AlphaFold2, AlphaFold3, and ESMFold with MCC = 0.611), while only two specialized predictors (spot_disorder2: 0.698; spot_disorder1: 0.692)Hanson et al. (2019) achieved comparable performance. Traditional methods like fidpnn Hu et al. (2021)(0.390) and espritz_d Walsh et al. (2012)(0.397) showed substantially lower accuracy, highlighting the advantage of structure-based confidence metrics.

This superior performance extends to discriminative power. AlphaFold2 achieved an AUC of 0.91 and F-max of 0.78, with AlphaFold3 nearly matching these scores (AUC = 0.90, F-max = 0.78), while ESMFold showed slightly lower but still strong performance (AUC = 0.87, F-max = 0.73) (Figure 1). The ROC and Precision-Recall curves demonstrate that pLDDT-based methods maintain excellent sensitivity and specificity across thresholds, with high precision even at elevated recall, critical for practical applications. To assess generalizability, we examined RMSD between predicted and experimental disorder across protein categories (Figure 2). For the complete dataset, AlphaFold2 and AlphaFold3 achieved low RMSD values (0.37-0.38). Performance varied by disorder content: highly ordered proteins showed exceptional accuracy (RMSD 0.20-0.22), while highly disordered proteins exhibited increased deviation (RMSD 0.48-0.58), suggesting pLDDT methods underestimate extreme disorder. Nevertheless, their consistent performance across all categories establishes pLDDT as a reliable tool that rivals or exceeds dedicated sequence-based disorder predictors.

**Figure 1.**
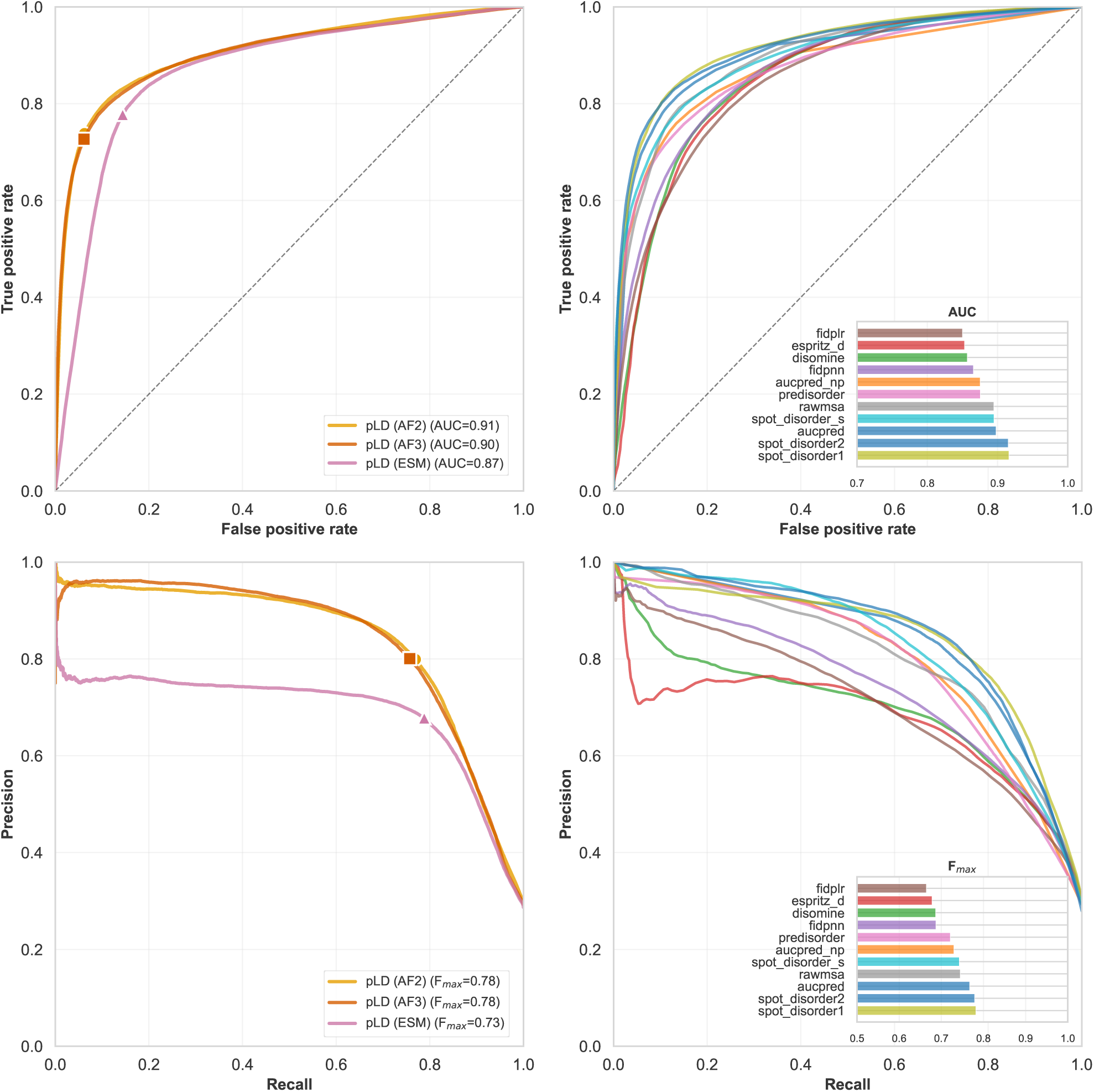
ROC and Precision-Recall analysis of disorder prediction performance. (Top left) ROC curves for pLDDT-based methods showing AUC values of 0.91 (AF2), 0.90 (AF3), and 0.87 (ESM). (Top right) Comparison of AUC scores across all methods tested. (Bottom left) Precision-Recall curves demonstrating F-max scores of 0.78 for both AF2 and AF3, and 0.73 for ESM. (Bottom right) F-max comparison across all methods. Squares on curves indicate optimal operating points. PLDDT-based methods consistently outperform specialized disorder predictors in both metrics.

**Figure 2.**
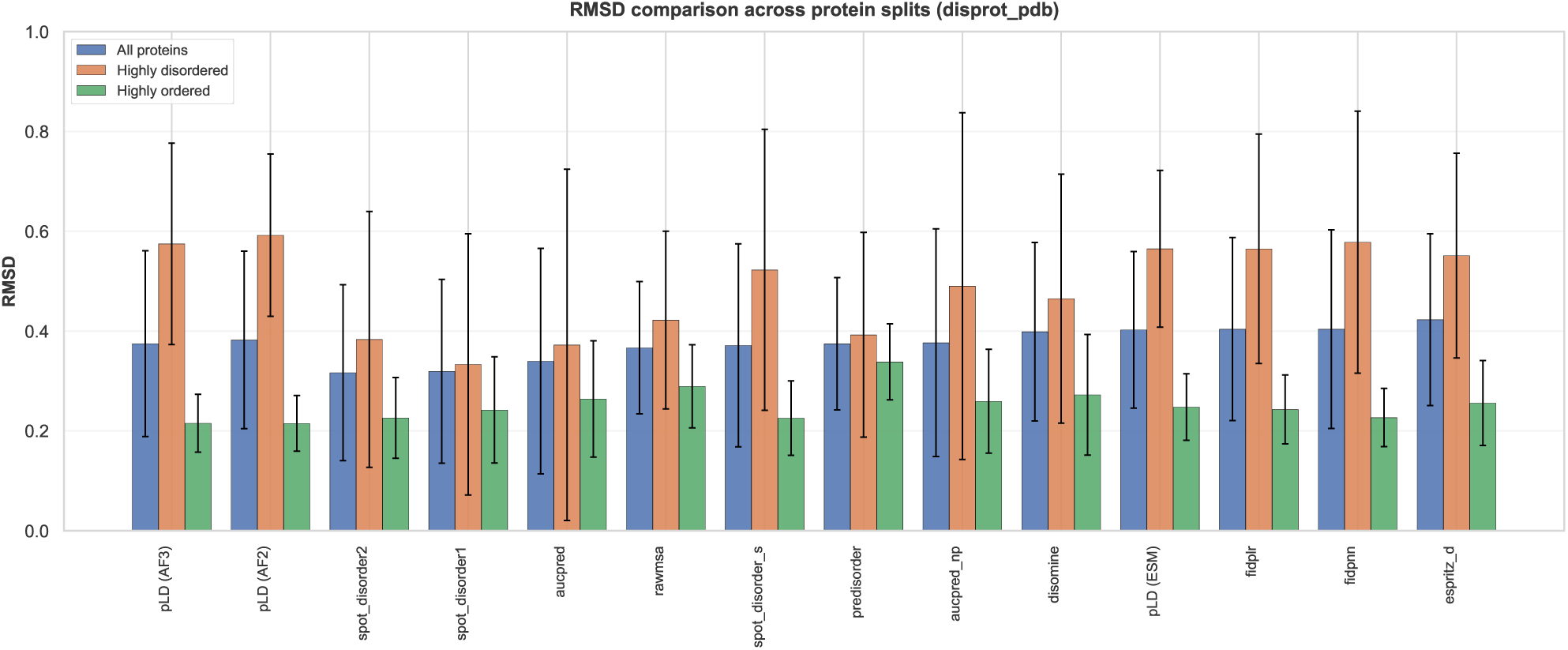
Root mean square deviation between predicted and experimental disorder across protein categories. RMSD values calculated for each predictor showing all proteins (left), highly disordered proteins with >50% disorder content (center), and highly ordered proteins with <10% disorder content (right). Error bars represent standard deviation. pLDDT-based methods show consistently low RMSD across categories, with particularly strong performance on highly ordered proteins with lower standard deviation.

### AlphaFold3 matches AlphaFold2 disorder prediction performance

While our benchmarking analysis demonstrated that AlphaFold2 slightly outperforms AlphaFold3 in overall metrics (MCC: 0.701 vs 0.693), a deeper examination of their prediction characteristics reveals remarkably similar performance profiles. To determine whether these models capture the underlying biophysical principles of protein disorder, we analyzed amino acid-specific disorder frequencies and distributions of per-protein prediction accuracy. Both AlphaFold2 and AlphaFold3 accurately recapitulate the amino acid disorder propensities observed in experimental data, achieving near-perfect correlations of r = 0.985 with ground truth frequencies (Figure S5). The models correctly identify disorder-promoting residues such as proline (P), serine (S), and glutamine (Q) with frequencies around 0.40, while accurately predicting low disorder propensity for hydrophobic residues like cysteine (C), isoleucine (I), and tryptophan (W) with frequencies below 0.20. ESMFold shows slightly lower but still strong correlation (r = 0.975), though it systematically overestimates disorder frequency across most amino acids, a trend most pronounced for disorder-promoting residues. This tight agreement with experimental amino acid preferences indicates that pLDDT-based predictions capture sequence-encoded disorder determinants rather than merely reflecting general structural confidence.

Analysis of per-protein prediction accuracy reveals complementary strengths between disorder and order recognition (Figure S3). For identifying disordered regions (true positive rate), AlphaFold2 and AlphaFold3 show similar performance distributions with medians of 0.779 and 0.762, respectively, though both exhibit substantial variation (means: 0.660 and 0.639). ESMFold demonstrates higher sensitivity to disorder (median: 0.848, mean: 0.723), consistent with its tendency to overpredict disorder. In contrast, all three models excel at identifying ordered regions (true negative rate), with AlphaFold2 and AlphaFold3 achieving exceptional accuracy (medians: 0.959 and 0.963; means: 0.903 and 0.908). ESMFold shows comparatively lower specificity for ordered regions (median: 0.899, mean: 0.798).

### Similar performances for AF2 and AF3 for multimers

To evaluate whether AF3’s architectural improvements enhance IDP-protein interaction modeling, we benchmarked AF2 and AF3 on 90 experimentally determined IDP-protein complexes. Despite AF3’s advances, both versions achieve remarkably similar performance across multiple structural and interface quality metrics. Both models showed nearly identical overall performance, with AF2 achieving marginally higher median DockQ scores (0.642 vs 0.625) and mean scores (0.571 vs 0.563) (Figure 3). This negligible 0.017 difference suggests AF3’s architectural changes neither substantially improve nor diminish IDP-complex prediction quality.

**Figure 3.**
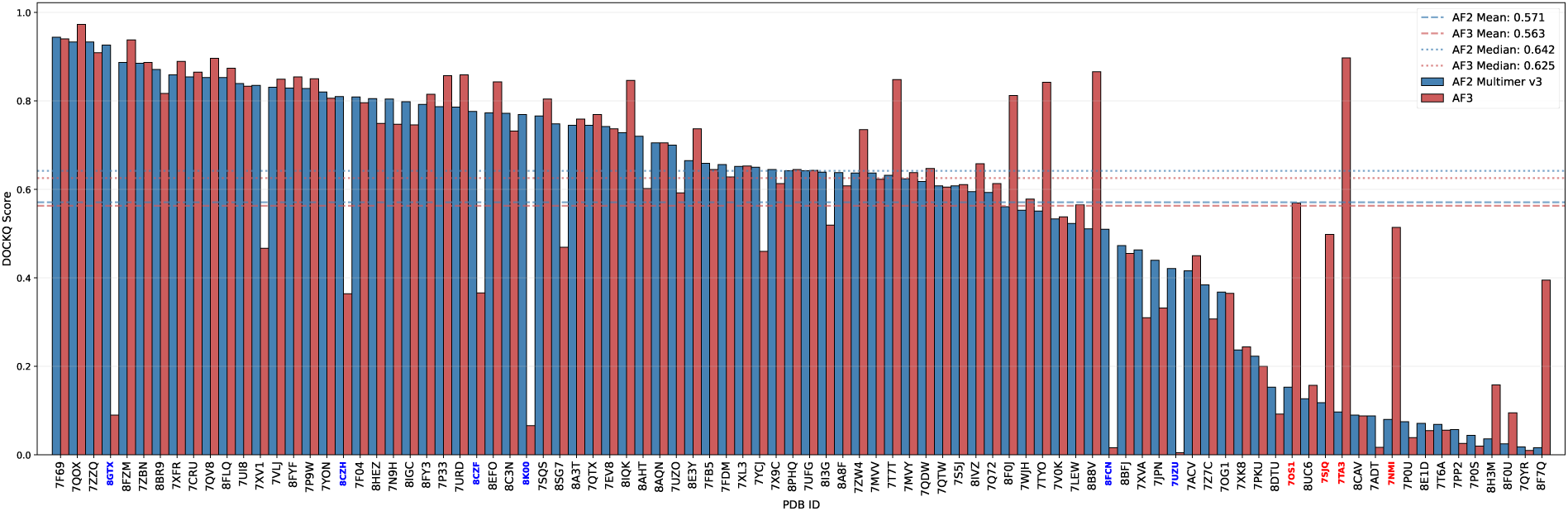
Overall structural quality comparison between AF2-Multimer and AF3. AlphaFold3 DockQ score distribution (mean: 0.563, median: 0.625, n=90). AlphaFold2-Multimer DockQ score distribution (mean: 0.571, median: 0.642, n=90). Both models achieve similar overall accuracy with marginal differences in central tendency. The top 10 cases with the largest DockQ score differences are highlighted with color-coded PDB IDs (blue: AF2-Multimer > AF3; red: AF3 > AF2-Multimer).

Interface contact analysis reveals similar challenges handling disordered regions (Figure S10). AF2 achieved a mean contact accuracy of 0.58 (54.4% over-contacted, mean disorder: 32.2%), while AF3 showed comparable performance (accuracy: 0.59, 51.1% over-contacted, mean disorder: 23.7%). Both exhibit a clear inverse relationship between disorder content and contact accuracy—complexes with high disorder (>40%) show substantially reduced accuracy, indicating neither model overcomes the challenge of predicting contacts in highly flexible regions. Performance equivalence extends across disorder levels (Figure S9). For low disorder interfaces (<20%), both achieve excellent mean DockQ (0.67), while high disorder interfaces (>40%) degrade similarly to 0.20 (mean AF3-AF2: −0.008). DSSP secondary structure matching correlates moderately with quality for both AF2 (r=0.524) and AF3 (r=0.487), suggesting accurate secondary structure prediction, though important, is not the primary determinant of overall quality.

Both models show similar dependencies on interface properties (Figure S11). Larger buried surface areas and higher hydrophobicity correlate with improved predictions, though effects are modest. AF2 shows minimal systematic effect from IDP coil content, while AF3 exhibits a slight performance reduction with high coil content. Secondary structure analysis reveals both predict predominantly helical conformations in IDP chains (median 0.60-0.62 helix, 0.05 sheet, 0.30-0.35 coil) (Figure S7)—a known limitation reflecting training on folded structures. Interface PAE distributions are nearly indistinguishable (Figure S6): AF2 mean 7.68 Å (median: 6.11 Å), AF3 mean 7.86 Å (median: 6.29 Å). Both show similar inverse PAE-DockQ correlations, suggesting comparable confidence calibration. CAPRI classification and success rates further confirm performance parity (Figure S8).

### Conventional Structural Features Explain Less of AF3’s DockQ Variance Than AF2’s

To understand the variation in multimer prediction quality between AF2 and AF3, we performed a retrospective regression analysis on the completed B90 predictions, asking how much of the already-observed DockQ variance, computed by comparing each model’s prediction to its known native structure, could be statistically explained by two conventional structural features extracted from the prediction itself in the form of interface disorder content and secondary structure matching (DSSP accuracy). We focused on two key metrics known to influence complex prediction accuracy, i.e., interface disorder content and secondary structure matching (DSSP accuracy). We first constructed univariate linear regression models with disorder fraction as the sole predictor of DockQ scores. Interface disorder showed the strongest correlation for both models, though with notable differences in predictive power (Figure 4, panels A and C). AF2 exhibited a strong negative correlation between disorder and prediction quality (*r* = *−*0.701, *R*^2^ = 0.492), indicating that a simple linear model using disorder content alone explains approximately 49% of DockQ variance. In contrast, AF3 showed a weaker relationship (*r* = *−*0.556, *R*^2^ = 0.310), with the linear model explaining only 31% of variance, a substantial reduction suggesting AF3’s performance depends less directly on disorder content.

**Figure 4.**
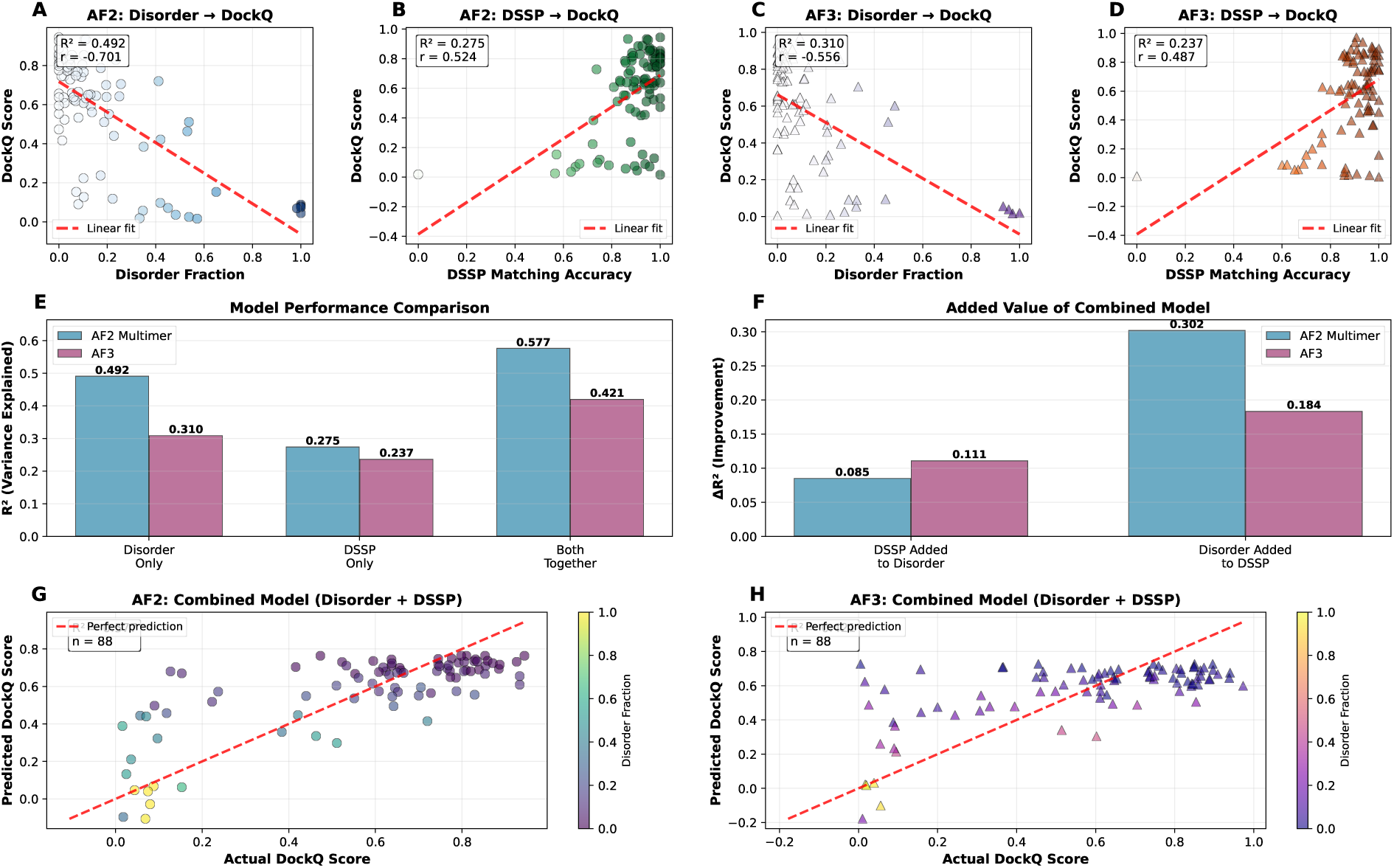
Regression analysis of AF2 and AF3 Dockq variations.

Linear regression models using secondary structure matching accuracy (DSSP) as the predictor demonstrated moderate positive correlations with DockQ for both models (Figure 4, panels B and D). AF2 achieved *r* = 0.524 (*R*^2^ = 0.275), while AF3 showed slightly lower correlation (*r* = 0.487, *R*^2^ = 0.237). This indicates that DSSP-based linear models account for only *∼*25% of prediction quality variance in both models.

For AF2, the combined linear model achieved *R*^2^ = 0.577, explaining 58% of DockQ variance (Figure 4, panels E and G). AF3 performed notably worse, with *R*^2^ = 0.421 explaining only 42% of variance (Figure 4, panel H). These multivariate models represent the maximum predictive power achievable using linear combinations of these two conventional metrics. Analysis of incremental *R*^2^ contributions (Figure 4, panel F) reveals asymmetric feature importance. For AF2, adding DSSP to a disorder-only linear model provided minimal improvement (Δ*R*^2^ = 0.085), while adding disorder to a DSSP-only model substantially improved prediction (Δ*R*^2^ = 0.302), indicating disorder is the dominant linear predictor for AF2 performance. In contrast, AF3 showed more balanced contributions (Δ*R*^2^ = 0.111 and 0.184), with neither feature dominating the linear model. The predicted versus observed DockQ plots (Figure 4, panels G and H) illustrate the linear models’ limitations. While both show general trends (points cluster near the diagonal), substantial scatter remains, particularly for AF3, where points deviate significantly from perfect prediction. Points are colored by disorder fraction, revealing that the largest prediction errors for AF3 occur across the full disorder spectrum, not just at high disorder levels, indicating that simple linear relationships between disorder and performance do not capture AF3’s behavior.

The finding that linear regression models using conventional structural metrics explain less than half of AF3’s DockQ variance and substantially less than for AF2 (42% vs 58%) has important implications, as this reduced predictability suggests AF3’s diffusion-based architecture introduces performance dependencies beyond standard interface properties and that these dependencies cannot be captured by simple linear relationships. The weaker linear correlation with disorder content (*R*^2^ = 0.310 vs 0.492 for AF2) indicates AF3’s failures are not simply explained by interface flexibility, as one might expect.

We emphasize that this analysis is diagnostic rather than predictive, as it does not estimate DockQ for novel, unsolved complexes, but instead probes whether a model’s prediction quality can be anticipated from basic properties of the prediction itself once the structure has already been generated and scored against experiment. The lower explanatory power for AF3 indicates that its successes and failures are harder to anticipate from these conventional features alone, which has practical implications for users deciding how much to trust a given AF3 output (see Discussion).

### AF3 Performance on MFIB

The inability to predict when AF3 succeeds or fails on general IDP complexes raises a critical question: how does AF3 perform on disorder-to-order transitions? To address this, we evaluated AF3’s performance on 90 complexes from the MFIB (Mutual Folding-Induced Binding) benchmark Fichó et al. (2025), where IDRs fold upon binding. AF3 shows comparable overall performance to AF2 on folding-induced binding. Analysis of prediction quality distributions reveals that both models achieve similar success rates across quality categories (Figure 5A). AF2 produced 42 excellent predictions (Interface RMSD *<* 1Å, Ligand RMSD *<* 2Å), 23 good predictions (I*<* 2Å, L*<* 3Å), 15 moderate predictions (I*<* 5Å, L*<* 5Å), and 10 poor predictions. AF3 showed remarkably similar performance with 43 excellent, 23 good, 16 moderate, and 8 poor predictions.

**Figure 5.**
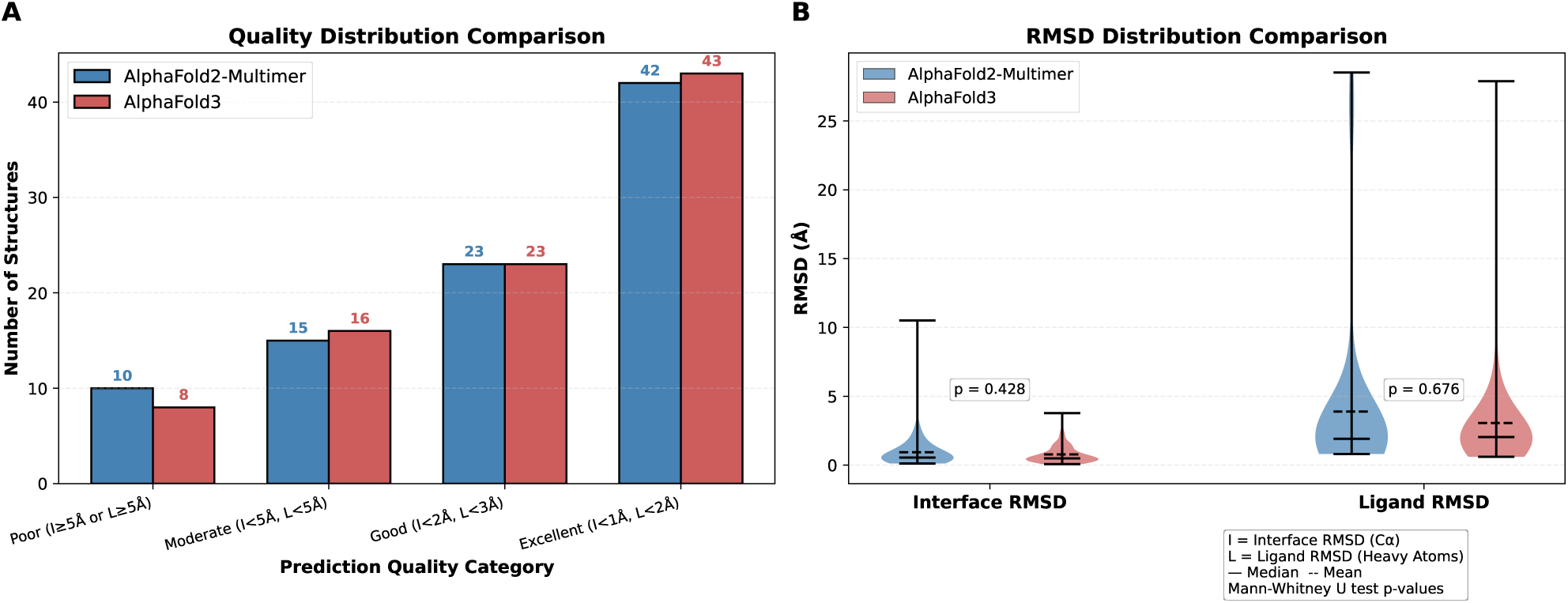
AlphaFold2-Multimer vs AlphaFold3 Performance on MFIB Benchmark Dataset. **(A)** Quality distribution comparison showing the number of structures in each prediction category: Poor (I*≥*5Å or L*≥*5Å), Moderate (I*<*5Å, L*<*5Å), Good (I*<*2Å, L*<*3Å), and Excellent (I*<*1Å, L*<*2Å). Both AlphaFold2-Multimer (blue) and AlphaFold3 (red) show similar distributions across quality categories for disorder-to-order transitions, with the majority of predictions achieving excellent quality. **(B)** RMSD distribution comparison showing violin plots for Interface RMSD (left) and Ligand RMSD (right). Both models show comparable median and mean RMSD values, with low Interface RMSD (median *∼*1Å) and slightly higher Ligand RMSD (median *∼*2.5Å). Horizontal lines indicate median (solid) and mean (dashed) values. I = Interface RMSD (C*α*); L = Ligand RMSD (Heavy Atoms). MFIB = Mutual Folding-Induced Binding benchmark.

The RMSD distribution analysis (Figure 5B) demonstrates that both models achieve comparable accuracy. Interface RMSD distributions show tight clustering around low values (median *∼*1Å) for both AF2 and AF3, while ligand RMSD distributions are similarly centered around 2.5Å. The overlapping distributions confirm near-equivalent performance on disorder-to-order transitions, suggesting that AF3’s diffusion-based architecture neither substantially improves nor worsens the prediction of complexes where IDRs fold upon binding compared to the established AF2 approach. The similar performance profiles indicate that both models successfully predict the final folded state of disorder-to-order transitions, likely because these complexes adopt well-defined structures in their bound forms—structures that are well-represented in PDB training data.

Additionally, we also examined AlphaFold performance on Molecular Recognition Features (MoRFs), short disordered segments that fold upon binding S12. Analysis reveals that both models struggle to identify these regions (Figure 6). AF2 shows greater confidence reduction in MoRF regions (mean Δ pLDDT = −2.07) versus AF3 (mean Δ = −0.96), yet direct prediction accuracy remains extremely low for both (AF2: 0.84%; AF3: 1.98% at optimal thresholds). MCW scores show no correlation with pLDDT values (AF2: r = −0.161, p = 0.129; AF3: r = −0.106, p = 0.322), confirming confidence metrics cannot identify MoRFs. This reveals a critical limitation: while both models successfully predict final folded states in the MFIB benchmark, they cannot prospectively identify which disordered regions will undergo binding-induced folding, limiting their utility for MoRF discovery.

### High-Divergence Cases Reveal Four Distinct Failure Modes

Next we seek to answer the question “Does AF3’s weaker disorder-dependence reflect genuine improvement in handling flexible interfaces, or a shift toward different failure modes?” We examined all complexes with substantial AF2–AF3 divergence (|ΔDockQ*| >* 0.3, *n* = 13), generating structural overlays for four representative cases (Figure S14). These cases resolve into four mechanistically distinct and internally consistent clusters as follows-

#### Cluster A, structured IDP (*n* = 5, AF2 wins all)

For low-disorder IDPs, AF2’s MSA-derived coevolutionary signal yields a tightly converged pair representation (interface PAE std: 2.2 Å in AF2 vs. 6.0 Å in AF3); Pairformer, relying on a reduced MSA signal compared to EvoFormer, produces noisier pair representations despite similar mean PAE. Cases: 8CZH, 8CZF, 8GTX, 7XV1, 8K00.^1^

#### Cluster B, diffusion collapse (*n* = 2, AF2 wins both)

AF3’s diffusion sampling fails to converge in the dominant prediction mode, producing uniform high uncertainty (PAE *>*25 Å, std *<*2 Å); AF2’s deterministic structure module is unaffected. Cases: 7UZU, 8FCN.^2^

#### Cluster C, genuine disorder (*n* = 3, AF3 wins all)

Where both models agree on high IDP disorder, AF3 achieves higher DockQ with lower PAE and higher confidence. In 7NMI, AF2 additionally misclassifies the receptor as disordered, collapsing the interface; AF3 preserves receptor order. Cases: 7NMI, 7OS1, 8F7Q.

#### Cluster D, MSA over-assignment (*n* = 3, AF3 wins all)

AF2’s MSA over-assigns disorder to IDPs that fold upon binding, likely from coevolutionary noise propagated from disordered homologs in the alignment; AF3, relying on a reduced MSA signal less susceptible to this contamination, correctly identifies the ordered or helical conformation. Cases: 7TA3, 8B8V, 7SJQ.

### Seed Variance Analysis Reveals Mostly Deterministic Failure with Rare, Case-Specific Exceptions

To determine whether AF3’s performance variation reflects stochastic sampling or systematic failure, we generated 30 additional AF3 predictions for six high-divergence cases (Figure S13; Table S4). Two cases (7TA3, 8K00) show tight clustering across all 30 seeds (std *<*0.04), confirming deterministic prediction whether correct (7TA3, mean 0.871, AF2 0.097) or incorrect (8K00, mean 0.062, AF2 0.769).

The remaining four cases are better characterized by distribution shape than summary statistics. Three (8CZF, 8CZH, 8FCN) show bimodal distributions as a large low-scoring majority and a discrete high-scoring minority. For 8CZF, 5/30 seeds (17%) score 0.79–0.81, nearly matching AF2 (0.776), while 25/30 score 0.35–0.38.

For 8CZH, 2/30 seeds (7%) reach *∼*0.81, matching AF2 (0.810), against 28/30 at 0.35–0.39. For 8FCN, 2/30 seeds (7%) reach 0.74–0.79, exceeding AF2 (0.510), against 28/30 below 0.13. The fourth case, 8GTX, shows no high-scoring mode: 29/30 seeds cluster at 0.27–0.42 (mean 0.294), well short of AF2 (0.926), with a single low outlier (0.029). The main-text value (DockQ 0.090) is a low-end seed rather than an atypically poor draw from an otherwise capable model.

Together, these patterns indicate that AF3’s diffusion process mostly commits to a fixed outcome regardless of seed; where variance exists, it reflects a rare discrete success mode rather than productive ensemble exploration.

## Discussion

This study presents the first systematic evaluation of AF3’s performance on IDPs and disorder-mediated interactions. Although AF3 reliably predicts disorder in monomers, its multimer modeling reveals unexpected complexity that conventional structural features cannot fully capture. AF3’s pLDDT scores reliably predict disorder, with performance comparable to AF2 (MCC: 0.693 vs 0.701) and rivaling specialized predictors. The near-perfect correlation in amino acid disorder propensities (*r* = 0.985) confirms that disorder prediction emerges from training data composition rather than architectural design, a phenomenon preserved across AF2’s Evoformer and AF3’s diffusion-based generation.

Despite similar aggregate performance (mean DockQ: 0.563 vs 0.571), AF2 and AF3 arrive at these results through different mechanisms. Interface disorder and secondary structure explain 58% of AF2’s DockQ variance but only 42% of AF3’s. Structural analysis of high-divergence cases (Results, “High-Divergence Cases Reveal Four Distinct Failure Modes”, Table S5, Figure S14) shows this reflects a shift in failure modes rather than genuine improvement: AF2 fails when EvoFormer’s MSA cannot anchor an ordered conformation or over-assigns disorder from disordered homologs; AF3 fails when its reduced MSA signal cannot resolve order, or when diffusion collapses to a degenerate state. AF3 succeeds relative to AF2 specifically where MSA contamination misleads EvoFormer or where genuine flexibility favors ensemble sampling. Both models perform well and comparably on disorder-to-order transitions (MFIB benchmark; majority of predictions Excellent; interface RMSD median *∼*1 Å; Mann-Whitney *p* = 0.428; Figure 5), consistent with these complexes adopting ordered bound conformations well-represented in PDB training data, confirming that AF3’s architectural changes neither improve nor worsen prediction when disordered regions fold into defined structures upon binding. Seed variance analysis on six high-divergence cases (30 seeds each; Table S4) reveals three patterns: two cases converge deterministically, three show bimodal distributions where 7–17% of seeds match or exceed AF2 while the majority fail, and one shows uniformly moderate performance with no high-scoring seeds, indicating AF3’s diffusion process can occasionally locate a correct solution without reliably converging to it, distinct from productive conformational ensemble exploration.

AF3’s bimodal behavior, producing excellent or poor predictions with few intermediate cases, creates practical challenges. For ordered interfaces, confidence metrics provide reliable quality estimates, but for disordered interfaces, these may be overconfident. We recommend comparing predictions across multiple seeds and between AF2 and AF3, as large divergences indicate structural ambiguity that warrants experimental validation; the failure-mode patterns identified above (Results) can help diagnose which model is more likely to be reliable for a given case, for example by comparing each model’s predicted interface disorder fraction or checking AF3’s interface PAE for the collapse signature described above. AF3’s diffusion framework struggles with disorder, not from algorithmic deficiency but from inheriting the PDB’s bias toward ordered structures. Architectural innovation alone cannot solve this problem. Progress requires disorder-enriched training data, ensemble prediction objectives, and evaluation metrics that capture conformational heterogeneity. Current benchmarks predominantly assess ordered proteins, creating circularity where models train on, are tested on, and are optimized for ordered structures, while disordered systems remain underrepresented.

Our study has limitations, as our 90-complex benchmark is modest; we analyzed only top-ranked predictions, and our feature analysis may not capture all relevant factors. Future work should explore whether fine-tuning or conditioning could improve AF3’s disorder handling. AF3 has not solved the fundamental challenge of modeling IDPs. While reliable for monomer disorder prediction, its multimer modeling operates through incompletely understood principles with unpredictable performance on disorder-containing complexes. For practitioners, this demands careful validation and comparison with experimental data. For the field, progress requires disorder-enriched training datasets and ensemble sampling strategies, not merely novel architectures trained on ordered-structure-biased data.

## Supplementary Information

**Figure S1.**
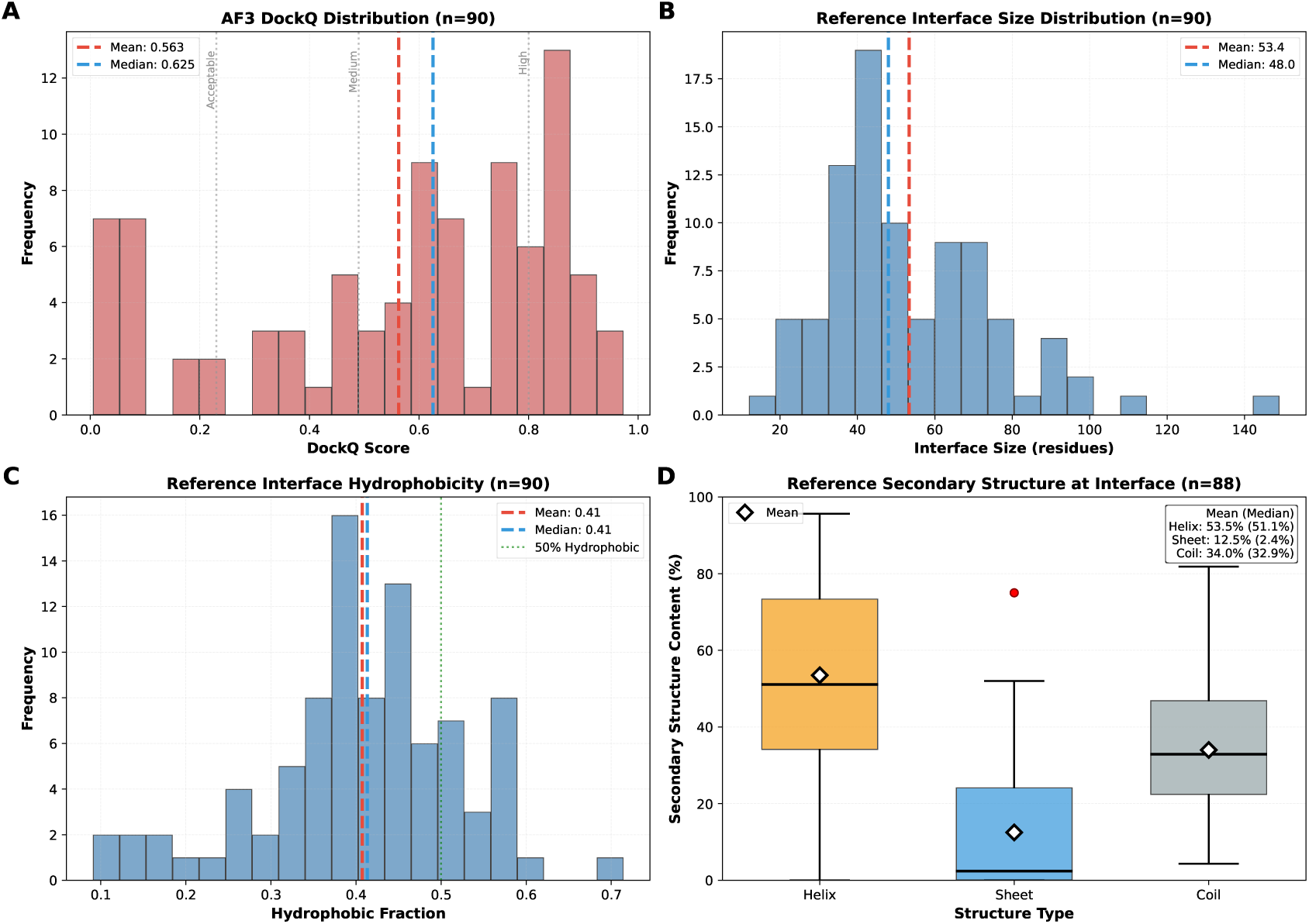
B90 dataset characterstics.

**Figure S2.**
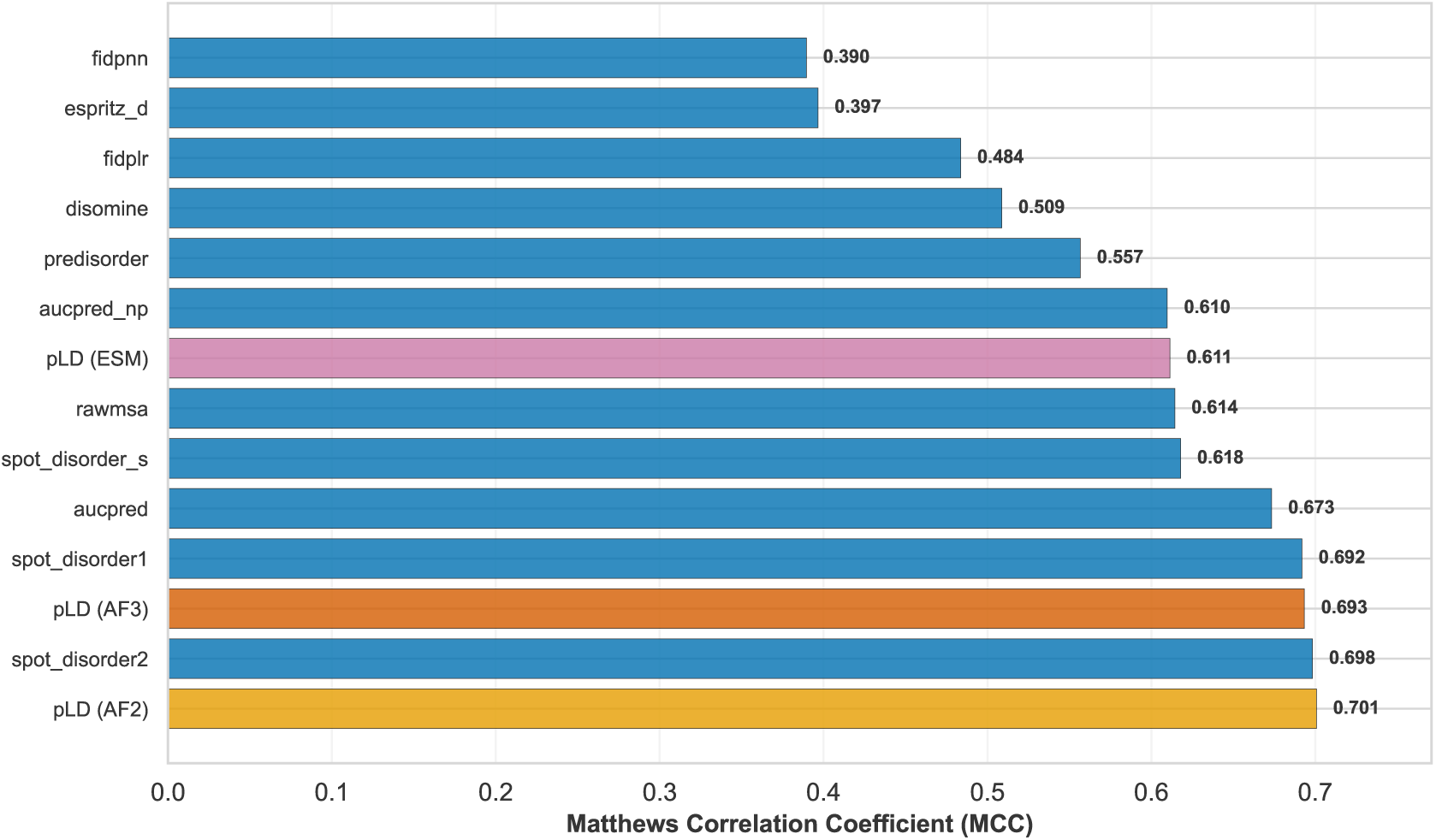
Matthews Correlation Coefficient comparison of disorder prediction methods. Benchmark results on the DisProt-PDB dataset showing MCC scores for pLDDT-based predictions from AlphaFold2 (AF2), AlphaFold3 (AF3), and ESMFold (ESM) compared to 11 sequence-based disorder predictors. pLDDT-based methods (highlighted in orange/pink) occupy the top three positions, with AlphaFold2 achieving the highest MCC of 0.701.

**Figure S3.**
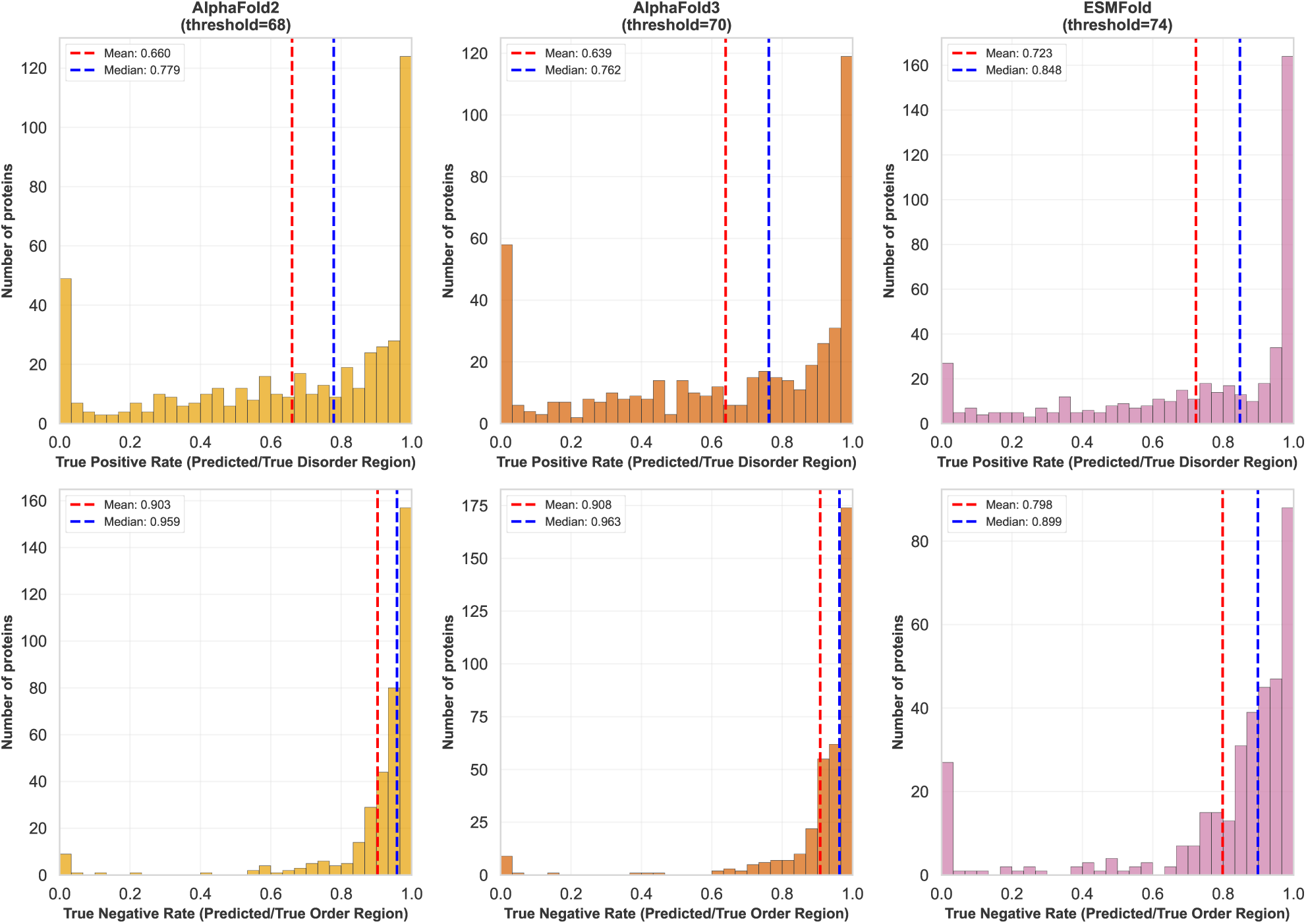
Distribution of per-protein disorder and order prediction accuracy. (Top row) True positive rate distributions showing sensitivity to disordered regions for AlphaFold2, AlphaFold3, and ESMFold at optimal thresholds (68, 70, and 74, respectively). (Bottom row) True negative rate distributions showing specificity to ordered regions. Dashed lines indicate mean (red) and median (blue) values. AlphaFold2 and AlphaFold3 show nearly identical profiles with exceptional specificity for ordered regions (medians >0.95) and moderate sensitivity to disorder. ESMFold exhibits higher disorder sensitivity but lower order specificity.

**Figure S4.**
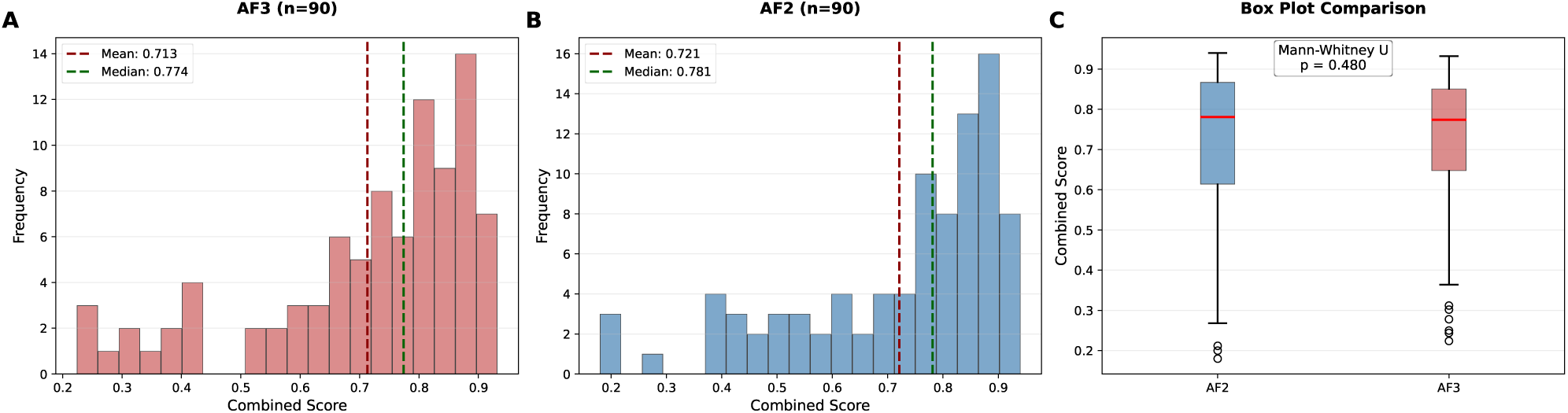
Combined confident score (0.8 x ipTM + 0.2 x pTM) comparison between AF2-Multimer and AF3. AlphaFold3 combined score distribution (mean: 0.713, median: 0.774, n=90). (B) AlphaFold2-Multimer combined score distribution (mean: 0.721, median: 0.781, n=90). (C) Box plot comparison showing nearly identical distributions with overlapping quartiles

**Figure S5.**
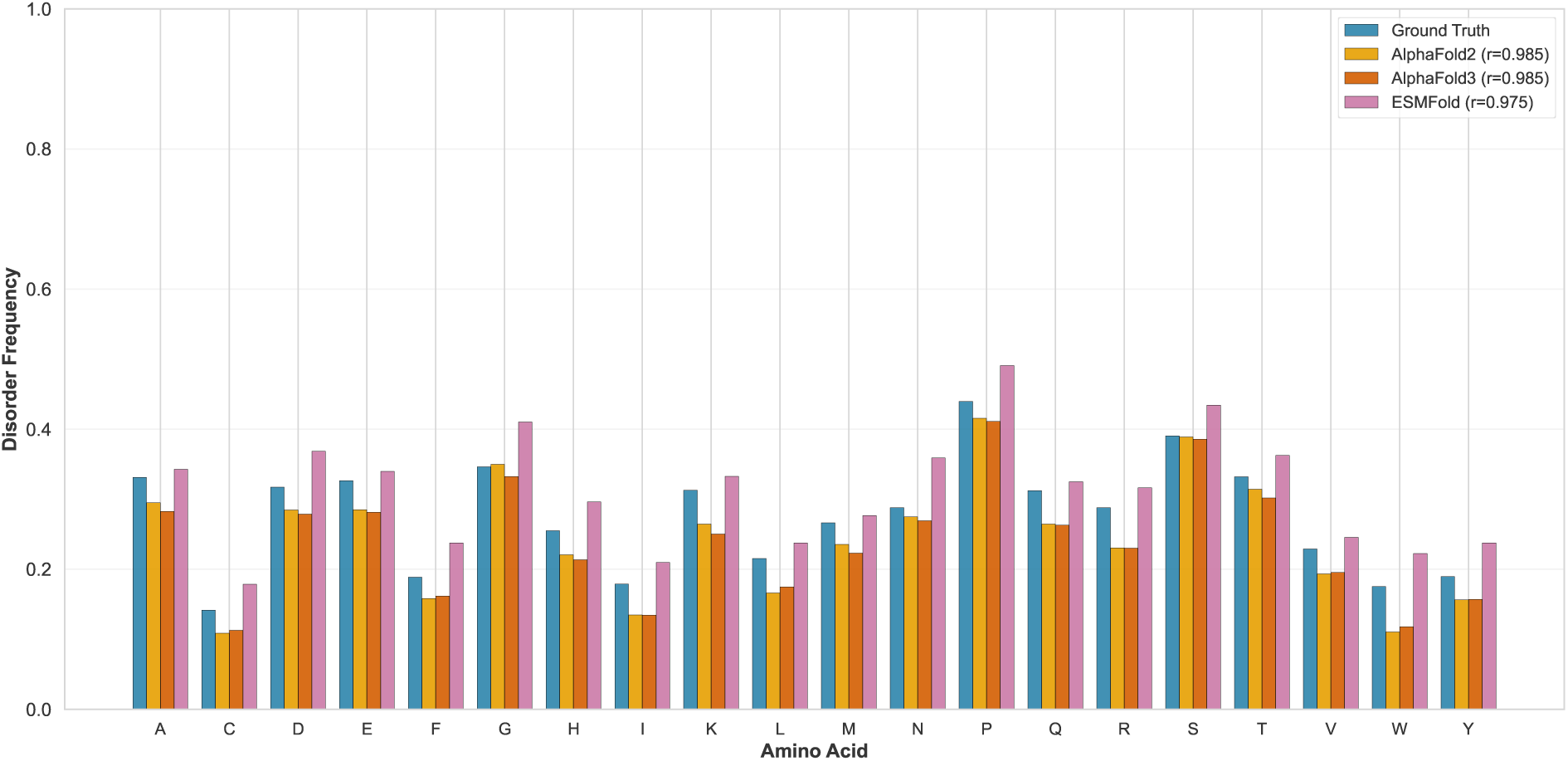
Amino acid disorder frequencies in ground truth and model predictions. Disorder propensity across all 20 amino acids comparing experimental annotations (blue) with pLDDT-based predictions from AlphaFold2 (yellow), AlphaFold3 (orange), and ESMFold (pink). Correlation coefficients with ground truth are shown in parentheses. Both AlphaFold2 and AlphaFold3 achieve r = 0.985, accurately capturing the spectrum from disorder-promoting residues (P, S, Q) to order-promoting residues (C, I, W). ESMFold (r = 0.975) shows systematic overestimation of disorder frequency.

**Figure S6.**
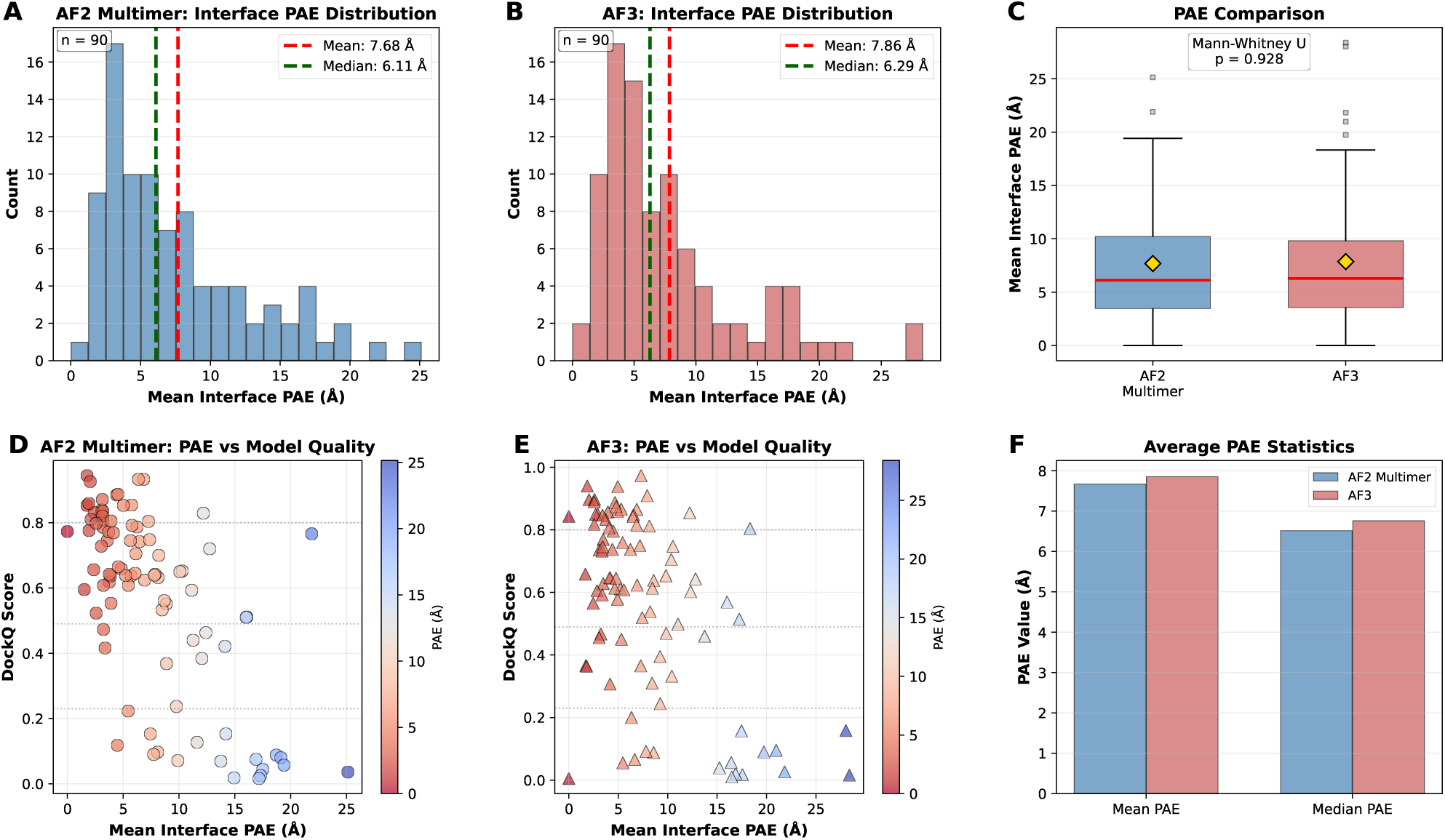
Predicted Aligned Error (PAE) analysis for interface confidence. (A-B) Interface PAE distributions for AF2-Multimer (mean: 7.68 Å, median: 6.11 Å) and AF3 (mean: 7.86 Å, median: 6.29 Å). (C) Box plot comparison showing nearly identical PAE ranges. (D-E) PAE versus DockQ scatter plots demonstrating inverse correlations for both models—low PAE indicates high-quality predictions. (F) Average PAE statistics confirm marginal differences. Both models provide similarly calibrated confidence estimates.

**Figure S7.**
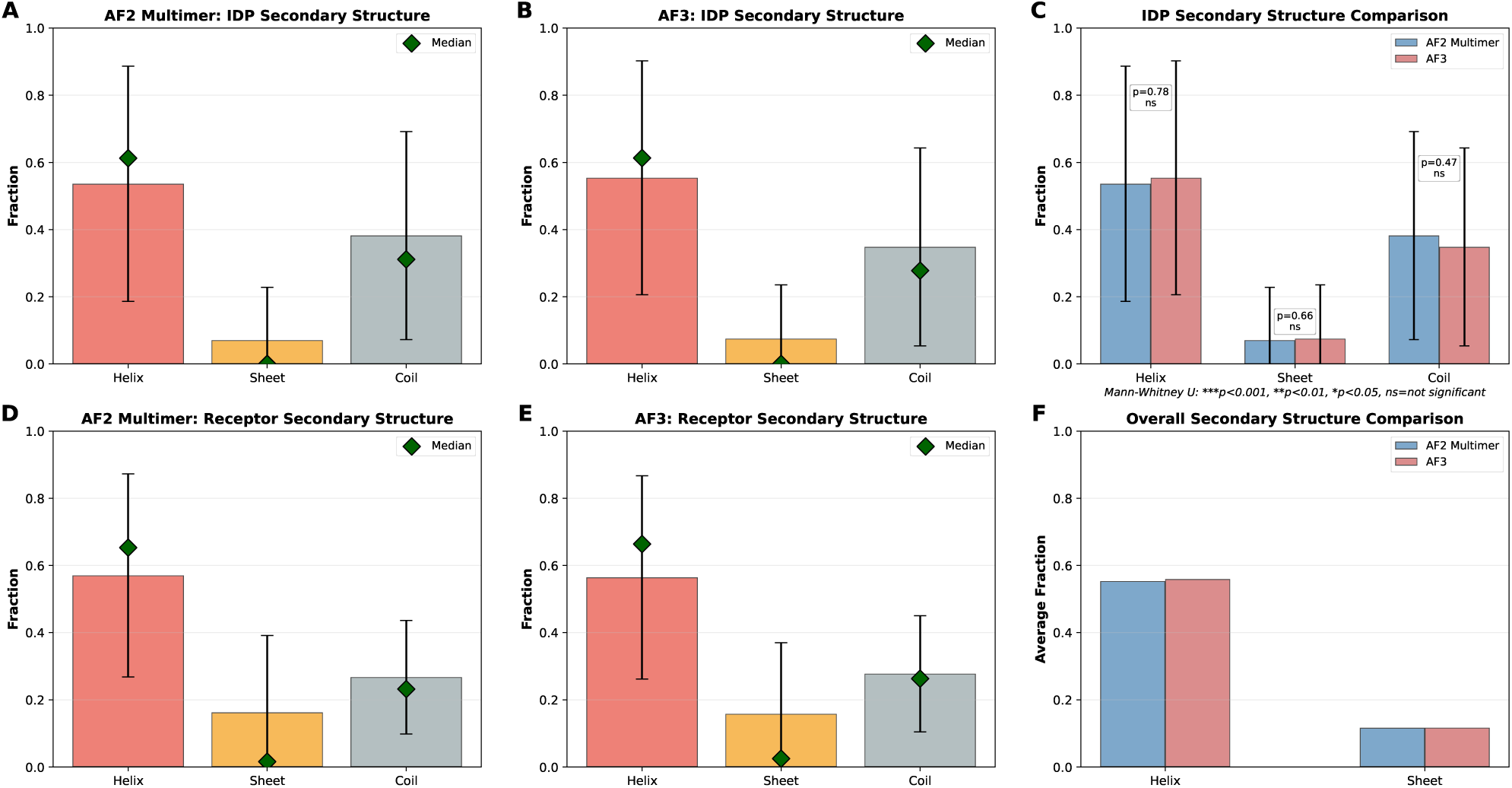
Secondary structure analysis of predicted complexes. (A-B) IDP chain secondary structure distributions for AF2-Multimer and AF3, both showing high helix content (median 0.60) and low sheet content. (C) Direct helix fraction comparison showing overlapping distributions. (D-E) Receptor secondary structures are nearly identical between models. (F) Overall secondary structure comparison confirms similar helix and sheet predictions. Both models tend to over-predict structured conformations in IDP regions.

**Figure S8.**
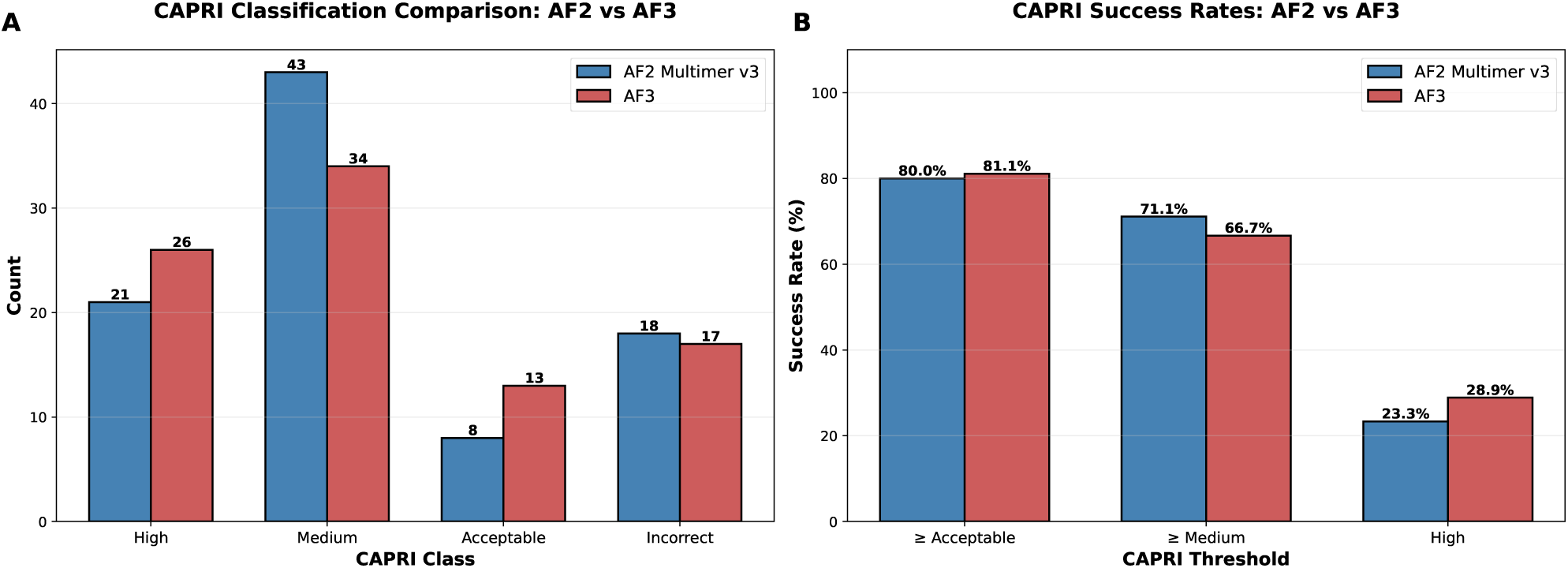
CAPRI divisions for AF2 and AF3.

**Figure S9.**
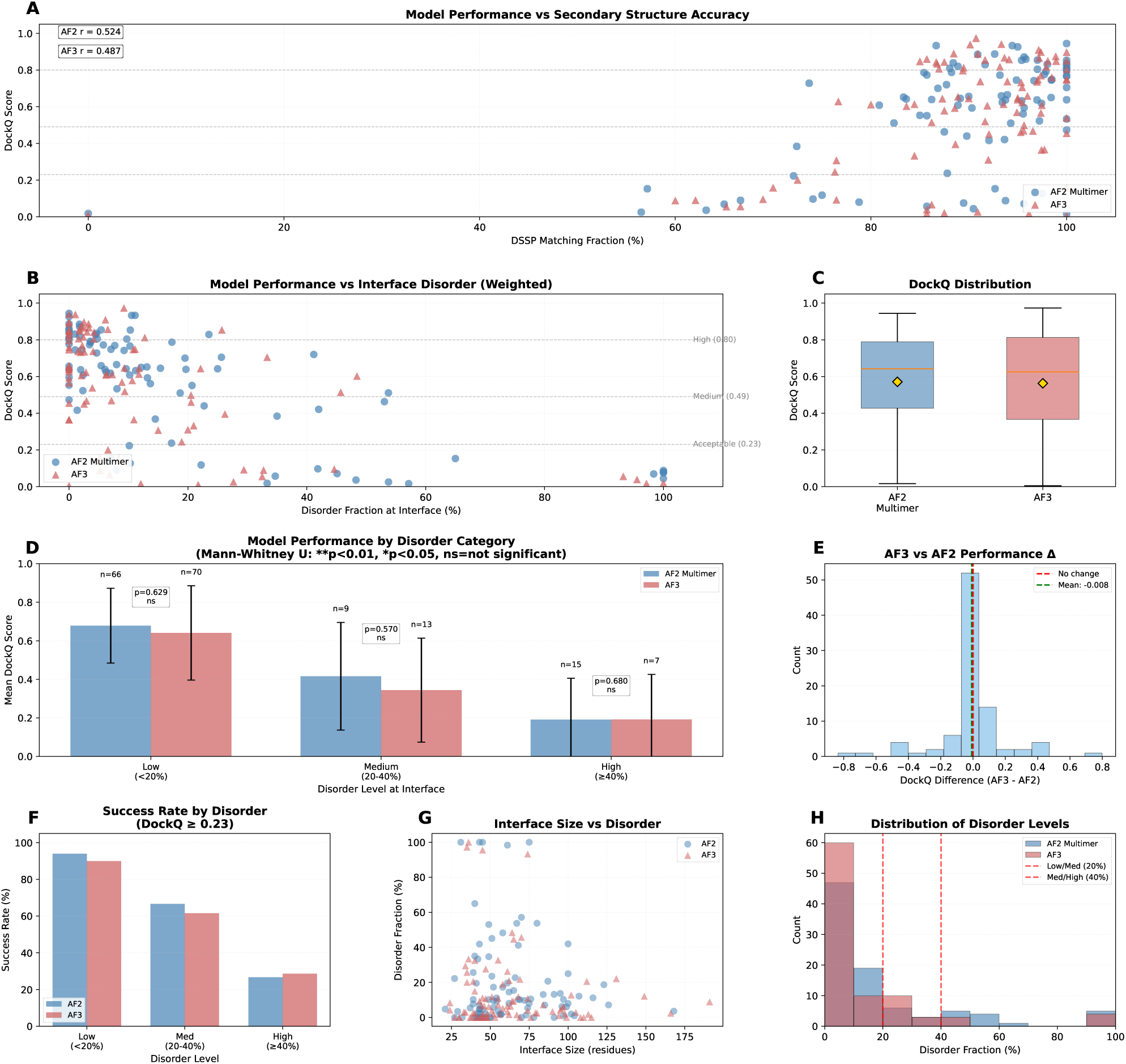
Comprehensive interface analysis comparing AF2-Multimer and AF3 performance. (A) DSSP secondary structure matching correlates moderately with DockQ (AF2: r=0.524, AF3: r=0.487). (B) DockQ versus interface disorder fraction showing similar degradation patterns. (C) Overall, DockQ distributions are nearly identical. (D) Performance by disorder category shows equivalent accuracy at low (*≤*20%), medium (20-40%), and high (>40%) disorder levels. (E) Performance difference histogram centered at −0.008. (F) Success rates (DockQ *≥*0.23) are comparable across disorder levels. (G) Interface size versus disorder shows no systematic relationship. (H) Disorder level distributions are similar between models.

**Figure S10.**
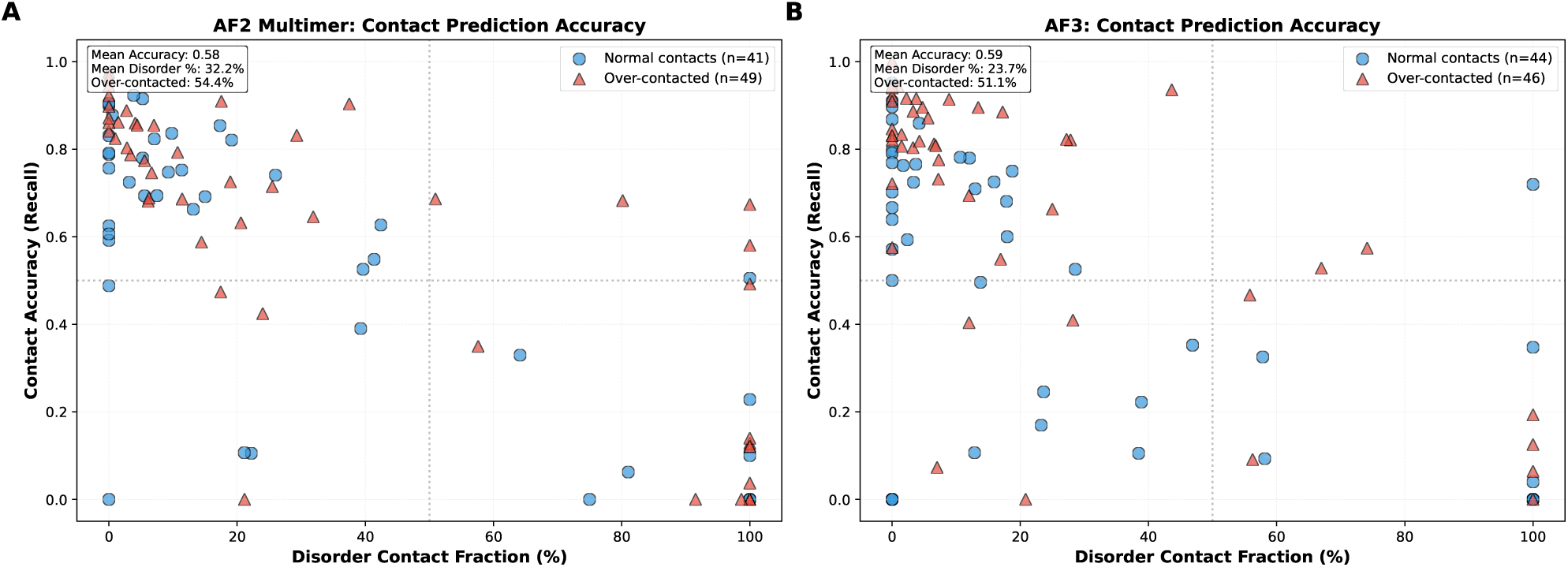
Interface contact prediction accuracy versus disorder content. (A) AlphaFold2-Multimer contact accuracy showing 54.4% over-contacted complexes with mean disorder of 32.2%. (B) AlphaFold3 showing 51.1% over-contacted complexes with a mean disorder of 23.7%. Both models show reduced contact accuracy at high disorder fractions, with normal contacts (blue circles) and over-contacted cases (red triangles) distributed similarly across disorder levels.

**Figure S11.**
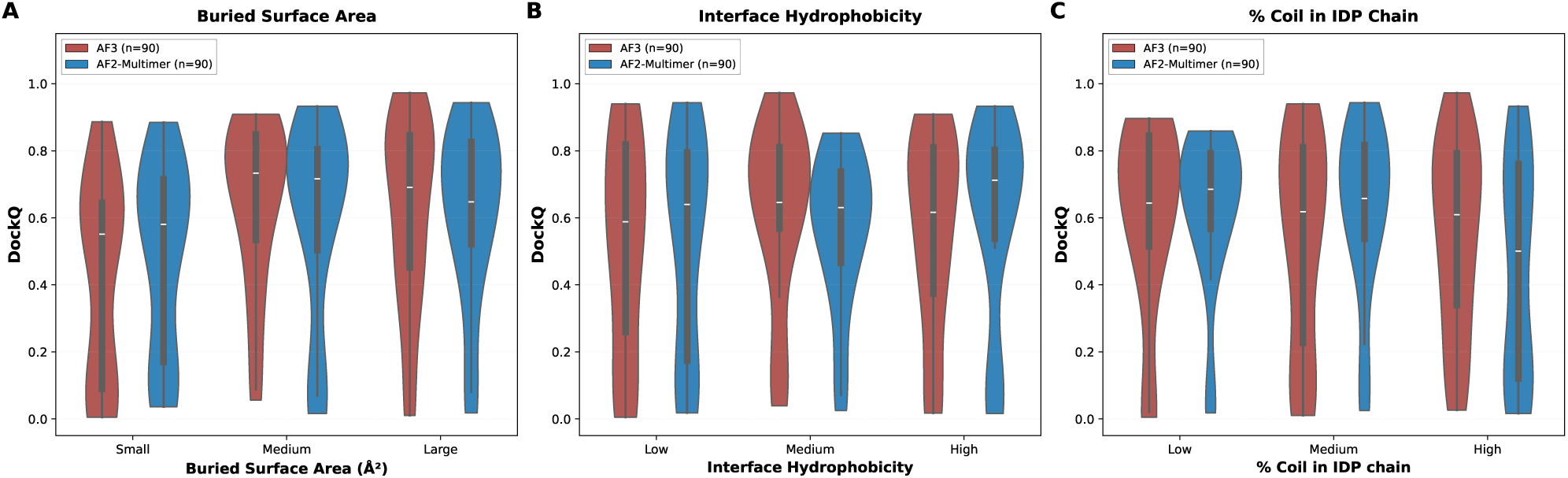
Structural determinants of model accuracy. (A-C) AlphaFold3 vs AlphaFold2 Multimer comparison for Buried Surface Area, Interface Hydrophobicity, and percentage of coil in the IDP Chain. Both models show similar dependencies: larger interfaces and higher hydrophobicity correlate weakly with improved accuracy, while coil content shows minimal systematic effect for AF2 and slight negative correlation for AF3. Violin plots show overlapping distributions across all categories.

**Figure S12.**
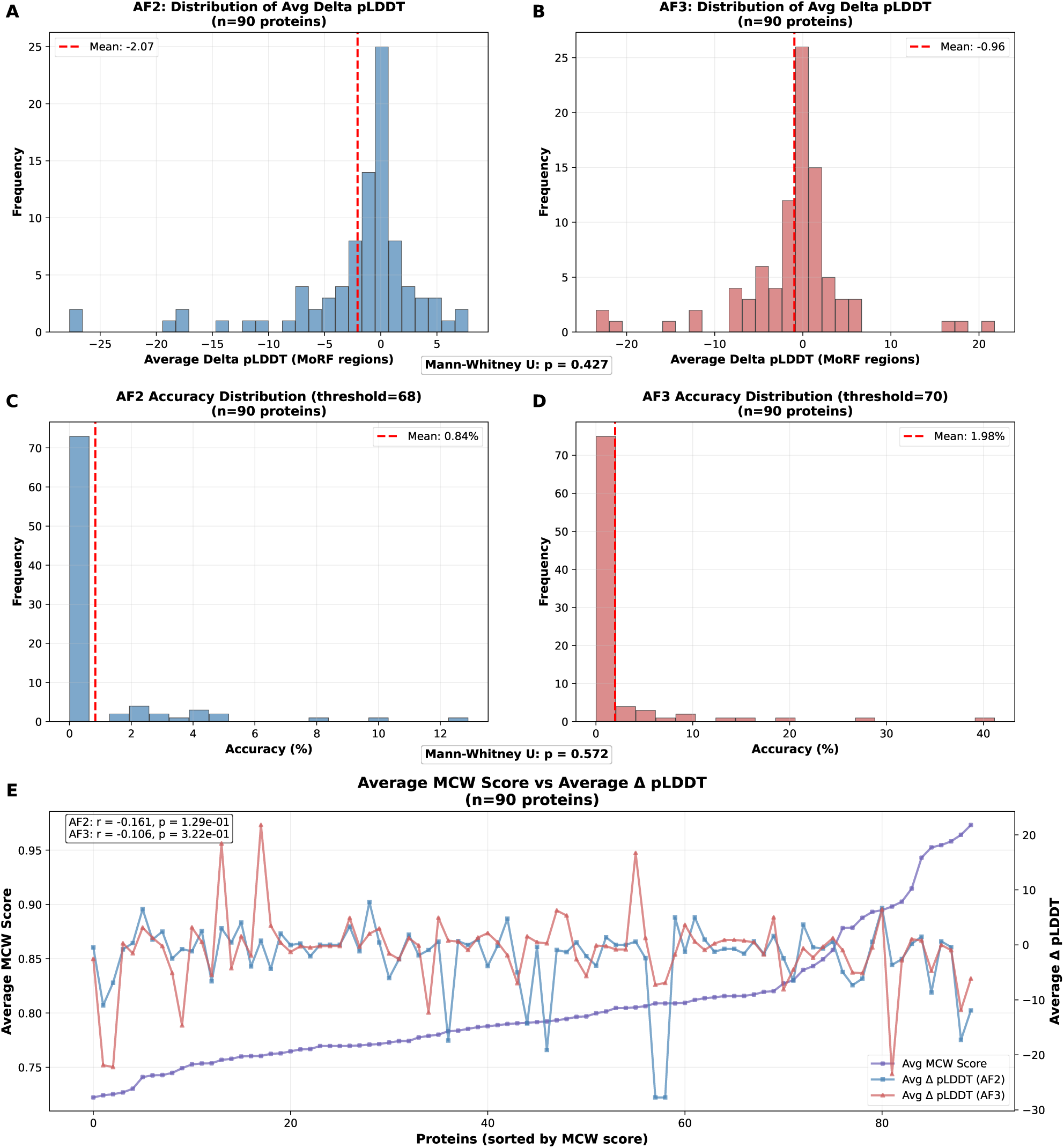
AlphaFold2 and AlphaFold3 struggle with Molecular Recognition Feature (MoRF) prediction. (A-B) Distribution of average delta pLDDT (difference between bound and unbound predictions) for MoRF regions across 90 proteins. AF2 shows greater confidence reduction (mean: −2.07) compared to AF3 (mean: −0.96). (C-D) Direct MoRF prediction accuracy distributions using optimal disorder thresholds (AF2: threshold=68, mean accuracy: 0.84%; AF3: threshold=70, mean accuracy: 1.98%). Both models show extremely low accuracy in identifying MoRF regions. (E) Relationship between average MCW (Mean Confidence Weighted) scores and average pLDDT values shows no significant correlation for either AF2 (r = −0.161, p = 0.129) or AF3 (r = −0.106, p = 0.322), indicating confidence metrics do not reliably distinguish MoRF regions. Despite successfully predicting final bound structures, both models fail to identify which disordered regions will undergo binding-induced folding prospectively.

**Figure S13.**
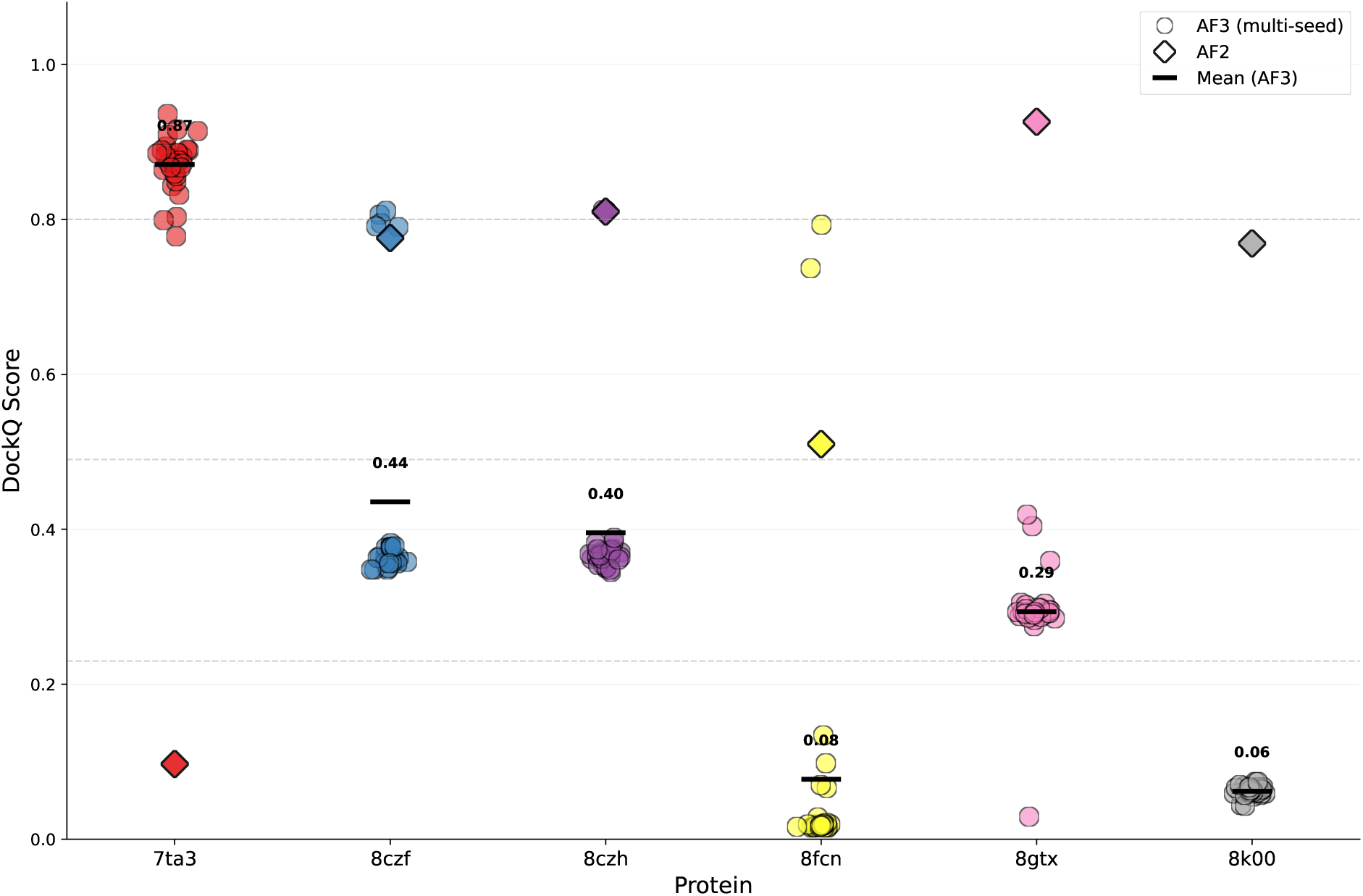
Multiseed analysis of AF2 and AF3 Dockq variations.

**Figure S14.**
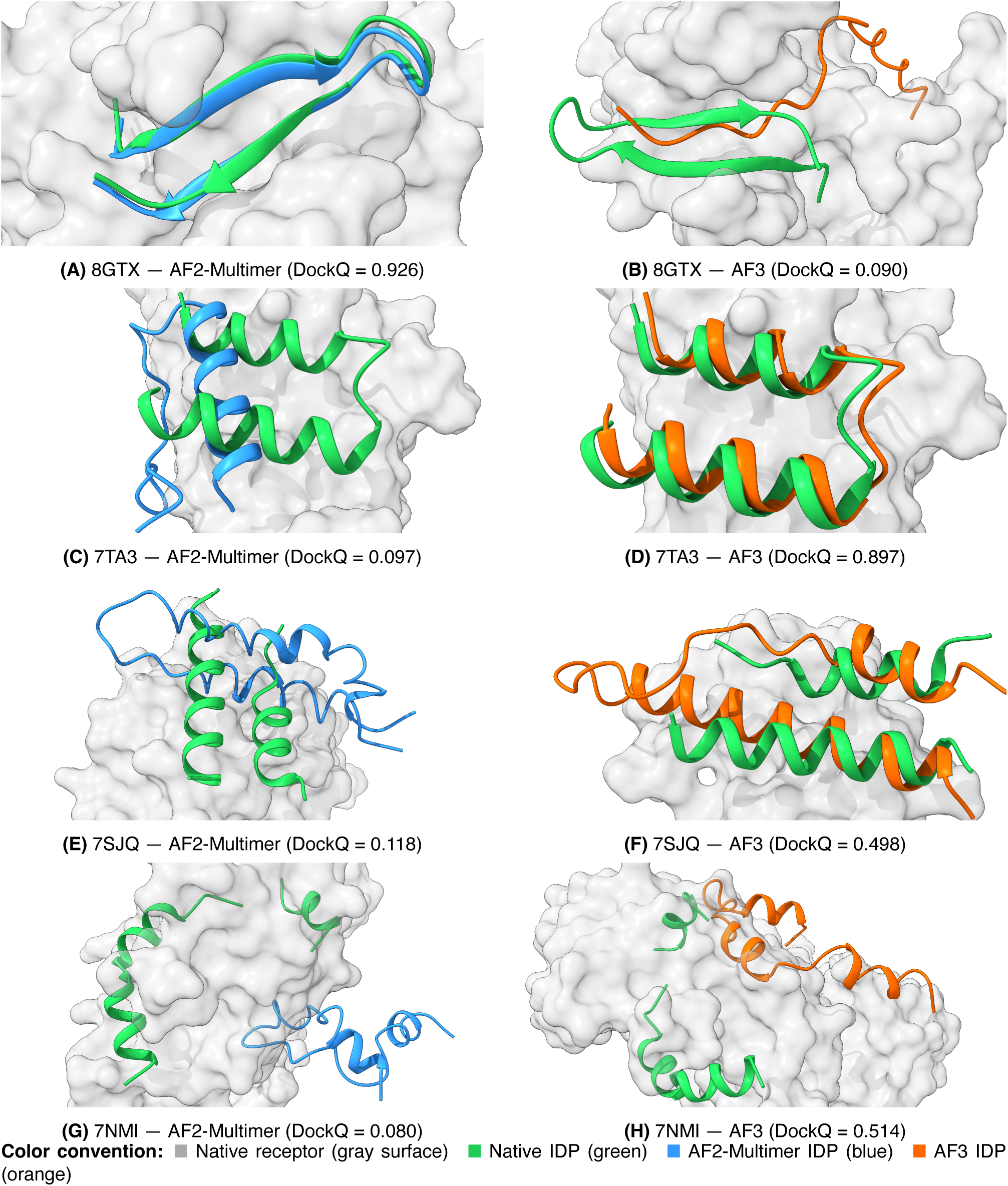
Structural overlays of four high-divergence cases. Predicted receptor chains were aligned to native (ChimeraX matchmaker) then removed; only predicted IDP chains shown. Retained only where alignment RMSD *<*2 Å on pruned pairs. **(A–B) 8GTX** (Spindlin1–HBX; |ΔDockQ| = 0.836). AF2 (DockQ 0.926) reproduces the native beta-hairpin via MSA-resolved order (disorder = 0.00); AF3 (DockQ 0.090) predicts full disorder (disorder = 1.00, PAE 19.6 Å) and misses the groove. **(C–D) 7TA3** (TNF*α*–cyclic peptide; |ΔDockQ| = 0.800; mirror of 8GTX). AF2 (DockQ 0.097) over-assigns disorder (MSA-contaminated, disorder = 0.79); AF3 (DockQ 0.897) correctly resolves the helix (helix = 0.74). **(E–F) 7SJQ** (PD-L1–cystine peptide; |ΔDockQ| = 0.380). AF2 (DockQ 0.118) predicts full disorder (helix = 0.40); AF3 (DockQ 0.498) identifies the helical MoRF (helix = 1.00). Slightly different viewing angles per panel. **(G–H) 7NMI** (S100P-ANXA2–p53; |ΔDockQ| = 0.434). IDP disordered in both models; failure mode is receptor handling. AF2 (DockQ 0.080) collapses the receptor (receptor disorder = 1.00); AF3 (DockQ 0.514) keeps it ordered (receptor disorder = 0.19). Slightly different viewing angles per panel.

## Supplementary Tables

**Table S1.**
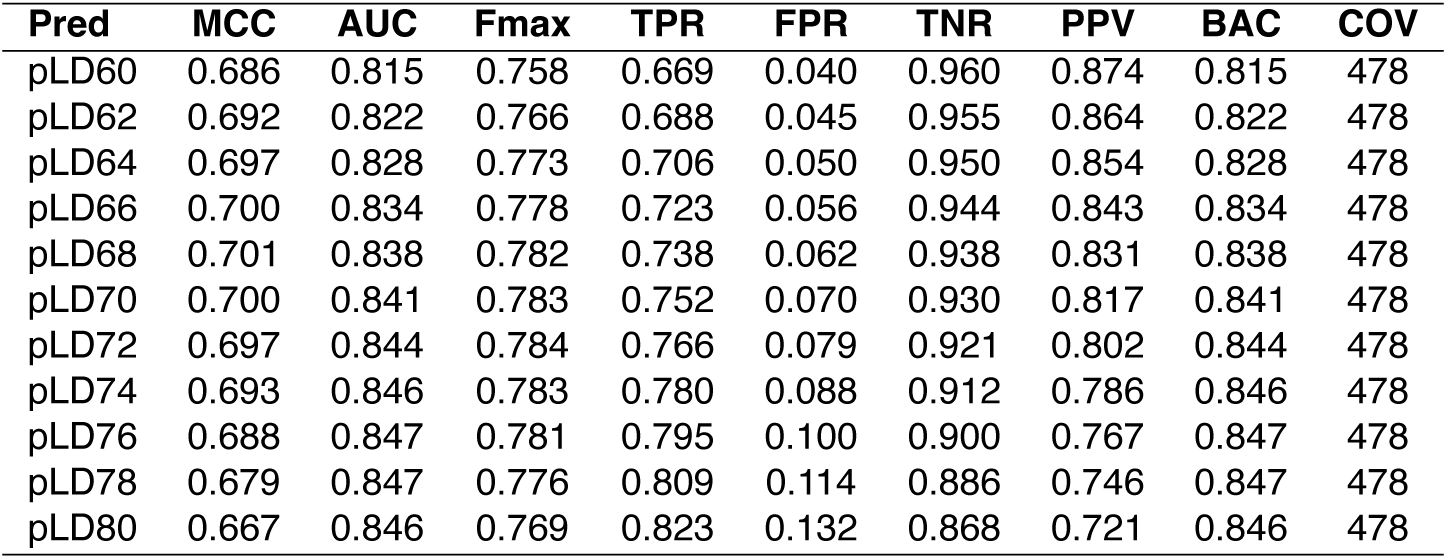
Performance metrics for AlphaFold2 predictions across pLD thresholds (60-80)

| Pred | MCC | AUC | Fmax | TPR | FPR | TNR | PPV | BAC | COV |
| --- | --- | --- | --- | --- | --- | --- | --- | --- | --- |
| pLD60 | 0.686 | 0.815 | 0.758 | 0.669 | 0.040 | 0.960 | 0.874 | 0.815 | 478 |
| pLD62 | 0.692 | 0.822 | 0.766 | 0.688 | 0.045 | 0.955 | 0.864 | 0.822 | 478 |
| pLD64 | 0.697 | 0.828 | 0.773 | 0.706 | 0.050 | 0.950 | 0.854 | 0.828 | 478 |
| pLD66 | 0.700 | 0.834 | 0.778 | 0.723 | 0.056 | 0.944 | 0.843 | 0.834 | 478 |
| pLD68 | 0.701 | 0.838 | 0.782 | 0.738 | 0.062 | 0.938 | 0.831 | 0.838 | 478 |
| pLD70 | 0.700 | 0.841 | 0.783 | 0.752 | 0.070 | 0.930 | 0.817 | 0.841 | 478 |
| pLD72 | 0.697 | 0.844 | 0.784 | 0.766 | 0.079 | 0.921 | 0.802 | 0.844 | 478 |
| pLD74 | 0.693 | 0.846 | 0.783 | 0.780 | 0.088 | 0.912 | 0.786 | 0.846 | 478 |
| pLD76 | 0.688 | 0.847 | 0.781 | 0.795 | 0.100 | 0.900 | 0.767 | 0.847 | 478 |
| pLD78 | 0.679 | 0.847 | 0.776 | 0.809 | 0.114 | 0.886 | 0.746 | 0.847 | 478 |
| pLD80 | 0.667 | 0.846 | 0.769 | 0.823 | 0.132 | 0.868 | 0.721 | 0.846 | 478 |

**Table S2.** Performance metrics for AlphaFold3 predictions across pLD thresholds (60-80)

| Pred | MCC | AUC | Fmax | TPR | FPR | TNR | PPV | BAC | COV |
| --- | --- | --- | --- | --- | --- | --- | --- | --- | --- |
| pLD60 | 0.678 | 0.806 | 0.747 | 0.647 | 0.035 | 0.965 | 0.883 | 0.806 | 478 |
| pLD62 | 0.683 | 0.812 | 0.754 | 0.663 | 0.039 | 0.961 | 0.875 | 0.812 | 478 |
| pLD64 | 0.687 | 0.818 | 0.761 | 0.679 | 0.044 | 0.956 | 0.865 | 0.818 | 478 |
| pLD66 | 0.690 | 0.823 | 0.767 | 0.696 | 0.049 | 0.951 | 0.854 | 0.823 | 478 |
| pLD68 | 0.693 | 0.828 | 0.772 | 0.712 | 0.055 | 0.945 | 0.842 | 0.828 | 478 |
| pLD70 | 0.693 | 0.833 | 0.775 | 0.727 | 0.061 | 0.939 | 0.831 | 0.833 | 478 |
| pLD72 | 0.692 | 0.836 | 0.777 | 0.740 | 0.069 | 0.931 | 0.817 | 0.836 | 478 |
| pLD74 | 0.690 | 0.839 | 0.778 | 0.755 | 0.077 | 0.923 | 0.802 | 0.839 | 478 |
| pLD76 | 0.687 | 0.841 | 0.778 | 0.771 | 0.088 | 0.912 | 0.785 | 0.841 | 478 |
| pLD78 | 0.680 | 0.843 | 0.775 | 0.787 | 0.101 | 0.899 | 0.763 | 0.843 | 478 |
| pLD80 | 0.668 | 0.842 | 0.769 | 0.803 | 0.119 | 0.881 | 0.737 | 0.842 | 478 |

**Table S3.** Performance metrics for ESMFold predictions across pLD thresholds (60-80)

| Pred | MCC | AUC | Fmax | TPR | FPR | TNR | PPV | BAC | COV |
| --- | --- | --- | --- | --- | --- | --- | --- | --- | --- |
| pLD60 | 0.528 | 0.743 | 0.640 | 0.569 | 0.083 | 0.917 | 0.733 | 0.743 | 468 |
| pLD62 | 0.547 | 0.757 | 0.662 | 0.605 | 0.090 | 0.910 | 0.729 | 0.757 | 468 |
| pLD64 | 0.563 | 0.771 | 0.679 | 0.638 | 0.097 | 0.903 | 0.725 | 0.771 | 468 |
| pLD66 | 0.579 | 0.783 | 0.695 | 0.671 | 0.104 | 0.896 | 0.721 | 0.783 | 468 |
| pLD68 | 0.591 | 0.794 | 0.707 | 0.700 | 0.113 | 0.887 | 0.715 | 0.794 | 468 |
| pLD70 | 0.602 | 0.803 | 0.717 | 0.728 | 0.121 | 0.879 | 0.707 | 0.803 | 468 |
| pLD72 | 0.608 | 0.811 | 0.724 | 0.753 | 0.132 | 0.868 | 0.697 | 0.811 | 468 |
| pLD74 | 0.611 | 0.817 | 0.728 | 0.778 | 0.144 | 0.856 | 0.685 | 0.817 | 468 |
| pLD76 | 0.609 | 0.820 | 0.728 | 0.799 | 0.160 | 0.840 | 0.668 | 0.820 | 468 |
| pLD78 | 0.602 | 0.820 | 0.724 | 0.820 | 0.179 | 0.821 | 0.648 | 0.820 | 468 |
| pLD80 | 0.592 | 0.818 | 0.716 | 0.840 | 0.203 | 0.797 | 0.625 | 0.818 | 468 |

**Table S4.** Multi-seed DockQ statistics for six cases exhibiting significant AF2–AF3 performance divergence. For each case, AF3 was run with thirty random seeds beyond the top-ranked prediction reported in the main analysis. Mean, standard deviation, minimum, and maximum DockQ are computed across all AF3 seeds; range is max minus min. AF2 DockQ (single prediction) is shown for comparison.

| Protein | Mean | Std | Min | Max | Range | AF2 DockQ |
| --- | --- | --- | --- | --- | --- | --- |
| 7TA3 | 0.871 | 0.034 | 0.778 | 0.936 | 0.158 | 0.097 |
| 8CZF | 0.435 | 0.165 | 0.348 | 0.811 | 0.463 | 0.776 |
| 8CZH | 0.396 | 0.114 | 0.345 | 0.812 | 0.467 | 0.810 |
| 8FCN | 0.077 | 0.189 | 0.015 | 0.793 | 0.778 | 0.510 |
| 8GTX | 0.294 | 0.060 | 0.029 | 0.419 | 0.390 | 0.926 |
| 8K00 | 0.062 | 0.007 | 0.043 | 0.074 | 0.031 | 0.769 |

**Table S5.** All 13 high-divergence cases (|ΔDockQ*| >* 0.3) with cluster assignments. Cases are grouped by mechanistic cluster (A–D) and sorted by |ΔDockQ| within each cluster. IDP disorder fraction is computed from predicted secondary structure; values near 0 indicate ordered, values near 1 indicate disordered prediction. Cluster A: structured IDP, EvoFormer advantage (AF2 wins all). Cluster B: diffusion collapse (AF2 wins both). Cluster C: genuine disorder, diffusion advantage (AF3 wins all). Cluster D: MSA over-assignment (AF3 wins all).

| Cluster | PDB | Winner | AF2 DockQ | AF3 DockQ | $\vert \Delta\text{DockQ}\vert $ | IDP Disorder Fraction | |
| --- | --- | --- | --- | --- | --- | --- | --- |
|  |  |  |  |  |  | AF2 | AF3 |
| A | 8GTX | AF2 | 0.926 | 0.090 | 0.836 | 0.00 | 1.00 |
| A | 8K00 | AF2 | 0.769 | 0.066 | 0.703 | 0.28 | 0.44 |
| A | 8CZH | AF2 | 0.810 | 0.364 | 0.446 | 0.00 | 0.00 |
| A | 8CZF | AF2 | 0.776 | 0.366 | 0.410 | 0.00 | 0.00 |
| A | 7XV1 | AF2 | 0.835 | 0.467 | 0.368 | 0.13 | 0.31 |
| B | 8FCN | AF2 | 0.510 | 0.016 | 0.494 | 1.00 | 1.00 |
| B | 7UZU | AF2 | 0.421 | 0.005 | 0.416 | 0.83 | 1.00 |
| C | 7NMI | AF3 | 0.080 | 0.514 | 0.434 | 1.00 | 1.00 |
| C | 7OS1 | AF3 | 0.153 | 0.569 | 0.416 | 0.86 | 0.73 |
| C | 8F7Q | AF3 | 0.016 | 0.395 | 0.379 | 1.00 | 0.82 |
| D | 7TA3 | AF3 | 0.097 | 0.897 | 0.800 | 0.79 | 0.00 |
| D | 7SJQ | AF3 | 0.118 | 0.498 | 0.380 | 1.00 | 0.50 |
| D | 8B8V | AF3 | 0.511 | 0.866 | 0.355 | 1.00 | 0.13 |

## Footnotes

1 Multi-seed analysis (30 seeds; Table S4) reveals bimodal behavior in two cases: 8CZF has 5/30 seeds (17%) at DockQ 0.79–0.81 against 25 seeds at 0.35–0.38; 8CZH has 2/30 seeds (7%) at *∼*0.81 against 28 seeds at 0.35–0.39. 8GTX shows no such high-scoring mode (29/30 seeds 0.27–0.42, mean 0.294; the main-text value of 0.090 is a low-end seed, not an atypically poor draw from an otherwise successful model). Seed means remain well below AF2 in all three cases.

2 8FCN shows a bimodal seed distribution (Table S4): 2/30 seeds (7%) reach DockQ 0.74–0.79, exceeding AF2 (0.510), while 28/30 cluster below 0.13. The top-ranked prediction (DockQ 0.016) falls in the collapsed mode.

## Notes

### Competing Interest Statement

The authors have declared no competing interest.

### Summary of Updates

The supplementary info, along with several important details, has been changed.

